# Nurse cell-derived small RNAs define paternal epigenetic inheritance in *Arabidopsis*

**DOI:** 10.1101/2021.01.25.428150

**Authors:** Jincheng Long, James Walker, Wenjing She, Billy Aldridge, Hongbo Gao, Samuel Deans, Xiaoqi Feng

**Author notes:** Jincheng Long, James Walker and Wenjing She contributed equally to this work.

## Abstract

The plant male germline undergoes DNA methylation reprogramming, which methylates genes *de novo* and thereby alters gene expression and facilitates meiosis. Why reprogramming is limited to the germline and how specific genes are chosen is unknown. Here, we demonstrate that genic methylation in the male germline, from meiocytes to sperm, is established by germline-specific siRNAs transcribed from transposons with imperfect sequence homology. These siRNAs are synthesized by meiocyte nurse cells (tapetum) via activity of the tapetum-specific chromatin remodeler CLASSY3. Remarkably, tapetal siRNAs govern germline methylation throughout the genome, including the inherited methylation patterns in sperm. Finally, we demonstrate that these nurse cell-derived siRNAs (niRNAs) silence germline transposons, thereby safeguarding genome integrity. Our results reveal that tapetal niRNAs are sufficient to reconstitute germline methylation patterns and drive extensive, functional methylation reprogramming analogous to piRNA-mediated reprogramming in animal germlines.

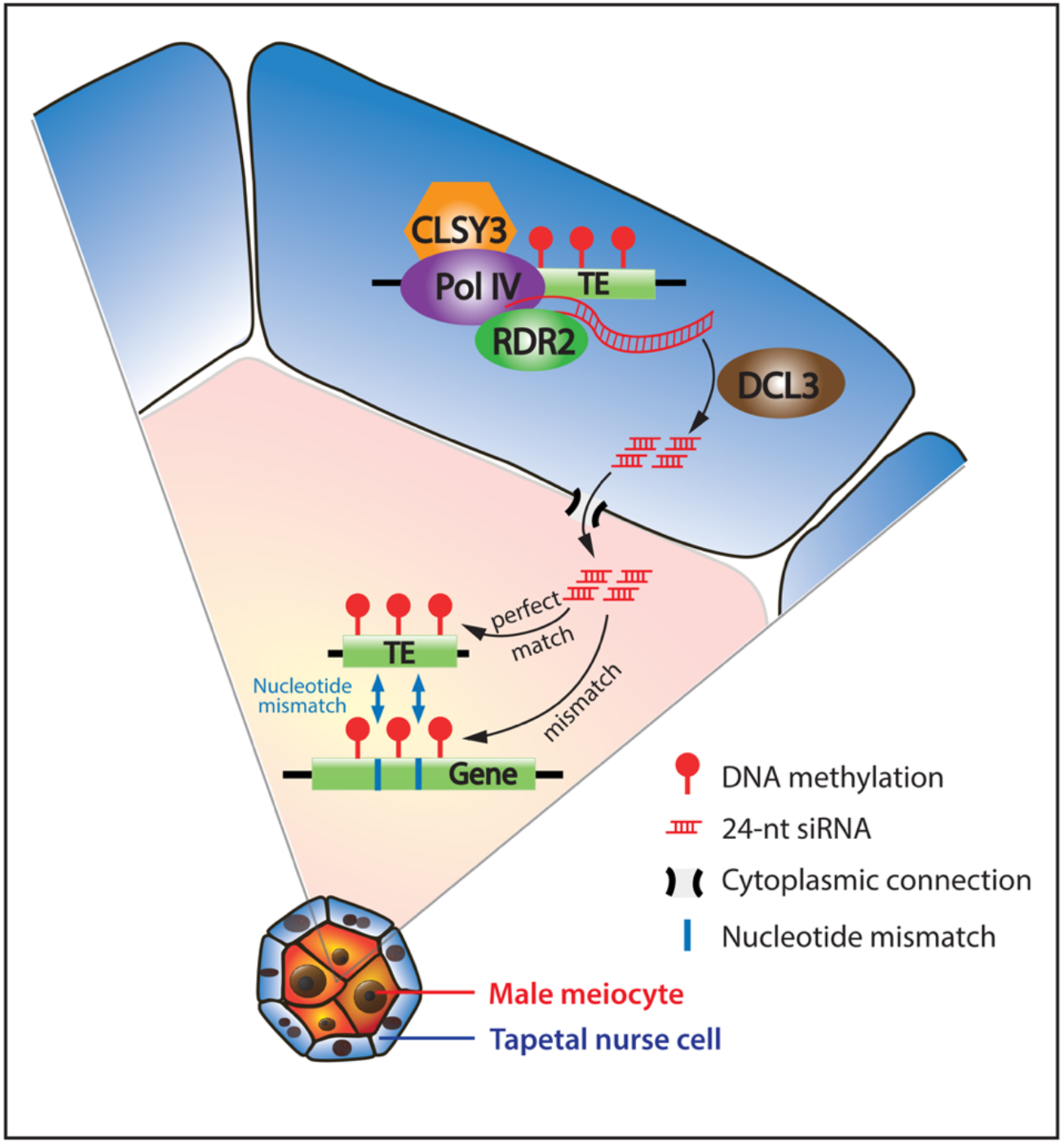

## Introduction

Methylation of the 5^th^ carbon of cytosine carries essential regulatory functions in eukaryotic genomes, including transcriptional regulation of genes and transposable elements (TEs)^1–3^. Cytosine methylation in the CG-dinucleotide context is maintained by DNA methyltransferase 1 (Dnmt1, called MET1 in plants), which methylates hemimethylated CG sites during DNA replication^4^. Plant TEs are also methylated in CHG and CHH contexts (where H is A, C, or T) by CMT3 (CHG) and CMT2 (CHH) methyltransferases^2^. Establishment of methylation *de novo* in all sequence contexts, and maintenance of CHG and CHH (non-CG) methylation, is catalyzed by plant Dnmt3 homologs (DRM1 and 2 in *Arabidopsis thaliana*)^4^. DRMs function within the small RNA-directed DNA methylation pathway (RdDM). In RdDM, 24-nt small interfering RNAs (siRNAs), which are produced from transcripts synthesized by RNA polymerase IV (Pol IV, a plant-specific derivative of Pol II) and RNA-dependent RNA polymerase 2 (RDR2), guide DRMs to target loci via association with a homologous transcript generated by Pol V (another plant-specific derivative of Pol II)^5^. Pol IV and Pol V preferentially associate with methylated DNA, making RdDM a self-reinforcing pathway, in which methylation promotes the generation of methylation-inducing siRNA^6,7^.

DNA methylation patterns are faithfully replicated during cell division, thus allowing methylation to have homeostatic functions during development^1,8^. However, in animals and plants, essential methylation reprogramming occurs in the germlines^9–11^. Mammalian germlines undergo genome-wide demethylation soon after their specification in the embryo^9^. This demethylation is crucial for epigenetic resetting, restoration of pluripotency and erasure of parental imprints^9,10^. Subsequently methylation is re-established globally, including imprints representative of the sex of the embryo^9^. Remethylation is mediated by Dnmt3 *de novo* methyltransferases and piwi-interacting RNAs (piRNAs), a class of siRNAs specifically expressed in gonads^12,13^. Impairment of methylation reprogramming reduces male and female fertility^13,14^. For example, male meiosis is arrested at the pachytene stage, associated with derepression of TEs and gene misregulation^13^.

Plants diverged from animals over a billion years ago from a common unicellular sexual ancestor^15^. As multicellularity evolved separately in plants and animals, they have adopted distinctive reproductive strategies^16,17^. Whilst animals usually have a reserved germline sequestered early in development, plants develop the germline from mature somatic cells^16,18^. Despite this difference, plants and animals convergently evolved specialized nurse cell lineages to nourish the developing germline^19^. For example, the male germline in *Arabidopsis* initiates as pollen mother cells (also called male meiocytes), which together with their surrounding nurse cells, tapetal cells (collectively called tapetum), descend from a common somatic precursor^19,20^ (Fig. 1a). Enclosed in this essential nurse cell layer, male meiocytes undergo meiosis to generate haploid microspores, each of which divides twice mitotically to give rise to two sperm and a companion vegetative cell in a pollen grain^17^ (Fig. 1a).

**Fig. 1:**
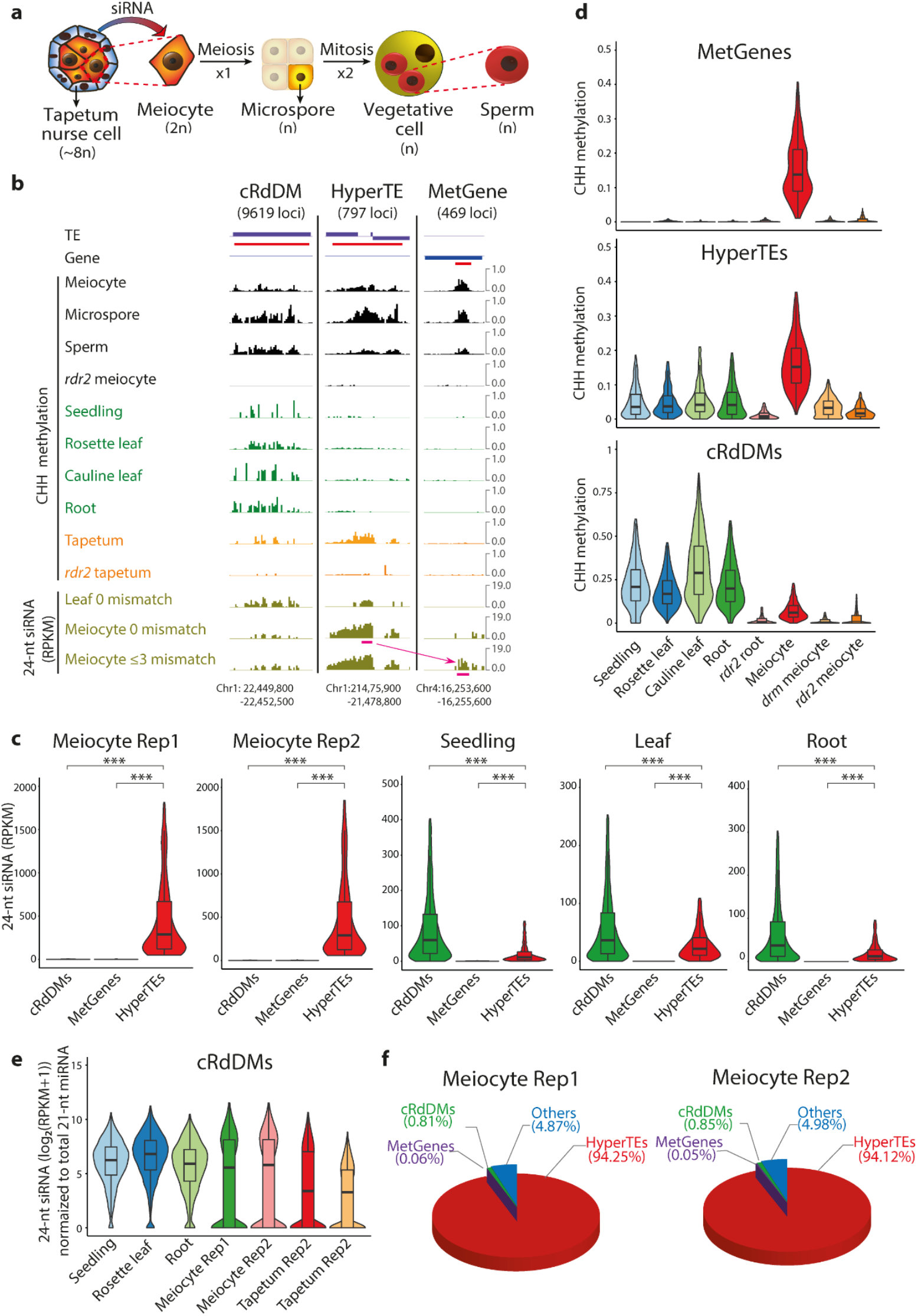
*Arabidopsis* meiocytes have a uniquely patterned small RNA transcriptome. **a**, Schematic diagram showing male germline development in *Arabidopsis*. n, number of chromosomes in the haploid genome. **b**, Snapshots of CHH methylation and 24-nt siRNA abundance at a canonical RdDM locus (cRdDM, left), a HyperTE (middle) and a MetGene (right) in different tissues and cells as labelled. Each category of locus is underlined in red, with the total number indicated in parentheses. 24-nt siRNAs in leaves and meiocytes are mapped with 0 or ≤3 mismatches as shown. siRNAs that target the MetGene with mismatches originate from the hyperTE, with both source and sink regions underlined in magenta. CHH methylation and siRNA (reads per kilobase per million; RPKM) levels are presented in 50-bp windows (w50s). For DNA methylation, w50s with at least ten informative cytosines are shown. **c**, Violin plots illustrating 24-nt siRNA levels at canonical RdDM loci (cRdDMs), HyperTEs and MetGenes in the male meiocyte, seedling, leaf and root. Note the different scales in the panels. Here, and in all subsequent violin plots, the plot shows the distribution of the data; the box encloses the middle 50% of the distribution (the horizontal line marks the median); and the whiskers illustrate 1.5 times the interquartile range. *** *P* < 2.2e-16, Kolmogorov-Smirnov test. Rep, biological replicate. **d**, Violin plots showing CHH methylation at MetGenes, HyperTEs and cRdDMs in somatic tissues and meiocytes. CHG and CG methylation are shown in Supplementary Fig. 1b. **e**, Violin plots depicting normalized 24-nt siRNA abundance against total 21-nt miRNAs in different somatic tissues, meiocytes and tapetal cells. **f**, Pie charts displaying percentages of 24-nt siRNAs associated with each category of locus in two meiocyte biological replicates.

Unlike mammals, plant male and female germlines do not undergo genome-wide demethylation and remethylation^11,21,22^. DNA demethylation does occur in gamete companion cells, the vegetative cell and its female equivalent^23–25^, but these lie outside the true germlines (Fig. 1a). The lack of drastic global reprogramming in germlines is consistent with the stable transgenerational transmission of methylation patterns observed in plants^21^. However, we recently characterized substantial functional methylation reprogramming in the *Arabidopsis* male germline. This reprogramming manifests most clearly at loci (mostly genes) that have non-CG methylation exclusively in the male germline (including meiocytes, microspores, vegetative and sperm cells), simplified as MetGenes (469 loci; Fig. 1b)^11^. This germline-specific methylation is important for meiosis progression, as the methylation at an essential meiotic gene facilitates correct splicing in meiocytes^11^. Germline methylation reprogramming is catalyzed by RdDM, and RdDM mutations such as *drm1;drm2* and *rdr2* affect meiosis and abolish non-CG methylation at MetGenes^11^. However, it remains unknown how genes become methylated by RdDM, a pathway that normally targets TEs, how particular genes are targeted, and how genic methylation specifically occurs in the germline.

Here, we show that genic methylation in male meiocytes is established by siRNAs transcribed from transposons with imperfect sequence homology. Production of these siRNAs occurs in the tapetal nurse cells and is mediated by the putative chromatin remodeler CLASSY3. The tapetal-produced siRNAs are transported into meiocytes and mediate methylation throughout the male germline, from meiocytes to sperm. We also demonstrate that tapetal siRNAs ensure germline genome integrity by suppressing TE expression. Altogether, our results reveal a class of nurse cell-derived siRNAs that induce methylation and regulate transcription in the entire male germline. The remarkable resemblance of tapetal siRNAs to animal piRNAs highlights the importance of specialized siRNA pathways for regulation of TEs and genes in germlines.

## Results

### Meiocyte siRNAs are distinctive from soma and mostly associate with a few TEs

To understand how specific loci are methylated by RdDM exclusively in the male germline, we sequenced siRNAs from isolated *Arabidopsis* male meiocytes (prophase I, mostly pachytene stage). In somatic tissues such as leaves, roots and seedlings, 24-nt siRNAs associate with methylated RdDM loci (~ 10,000 loci, overwhelmingly TEs; Fig. 1b,c)^11^. In relation to somatic tissues, nearly all (98%) of these canonical RdDM loci have many fewer 24-nt siRNAs in meiocytes (Fig. 1b,c and Supplementary Fig. 1a), and CHH methylation at these loci is greatly reduced (Fig. 1b,d). However, when normalized to 21-nt microRNAs, the levels of 24-nt siRNAs at canonical RdDM loci are similar in meiocytes and soma (Fig. 1e). The vast relative difference in 24-nt siRNA abundance occurs because male meiocyte 24-nt siRNAs are highly concentrated (log_2_(RPKM+1) > 7) in 854 clusters, most of which (80%) do not overlap canonical RdDM loci. These clusters have higher levels of siRNAs and non-CG methylation in meiocytes compared to somatic tissues (Fig. 1b,c,d and Supplementary Fig. 1b), and substantially overlap the 724 loci we previously identified as hypermethylated in the male germline^11^ (Supplementary Fig. 1c). Consistent with RdDM targeting, the overwhelming majority (93%, 797) of meiocyte siRNA clusters have significantly (*P* < 0.01, Fisher’s Exact test) reduced CHH methylation in *drm1 drm2* double mutant (simplified as *drm*) or *rdr2* mutant (Fig. 1b,d and Supplementary Table 1). These 797 siRNA clusters span 588 Kb (0.4% of the genome) but comprise 94% of the clustered 24-nt siRNAs, mostly (70%) overlapping TEs (TEs make up 18% of the *Arabidopsis* genome; Fig. 1f and Supplementary Table 2). Altogether, our results demonstrate that male meiocytes have a distinctive RdDM-associated siRNA profile, with the vast majority of 24-nt siRNAs targeting a small number of hypermethylated TEs (we will refer to these 797 loci as HyperTEs; Supplementary Table 2).

### HyperTE-derived siRNAs induce methylation at imperfectly matching meiocyte genes

Surprisingly, we found few 24-nt siRNAs in meiocytes associated with MetGenes (Fig. 1b,c,f), even though MetGenes are targeted by RdDM^11^. A careful comparison revealed that MetGenes and HyperTEs share similar sequences, leading us to hypothesize that MetGenes may be targeted by siRNAs produced from HyperTEs. Supporting this hypothesis, we found that 24-nt siRNAs generated from HyperTEs can be aligned to MetGenes if up to three mismatches are allowed (Figs. 1b and 2a and Supplementary Table 3). Importantly, the association of siRNAs with MetGenes is not caused by random mapping of mismatched siRNAs, because neither all genes nor random control loci associate with noticeable levels of mismatched siRNAs (Fig. 2a). Furthermore, we find that for 89% of HyperTE-associated siRNAs that map to MetGenes, MetGenes are the best hits in the genome aside from the source HyperTEs. These analyses suggest that siRNAs produced from HyperTEs cause methylation at MetGenes with similar sequences.

**Fig. 2:**
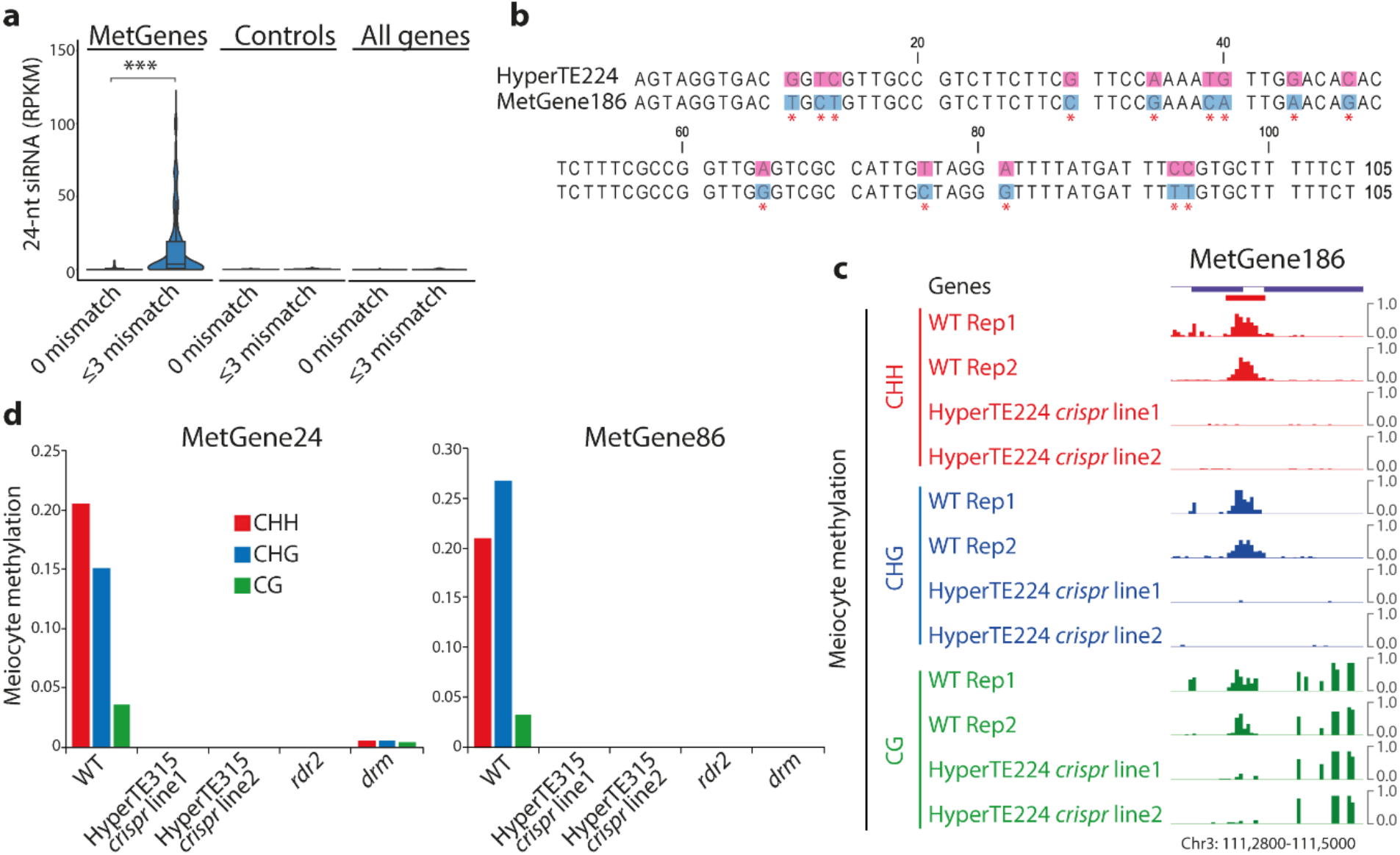
24-nt siRNAs produced from HyperTEs drive DNA methylation at MetGenes with mismatches. **a**, Violin plots illustrating the abundance of 24-nt siRNA mapped with 0 or ≤3 mismatches to MetGenes, a control set of random loci of similar sizes, and all genes in meiocytes. *** *P* < 2.2e- 6, Kolmogorov-Smirnov test. **b**, DNA sequence alignment between HyperTE224 (*AT2TE15980*) and the corresponding MetGene (MetGene186, *AT3G04230*), where HyperTE224-associated 24-nt siRNAs map with up to 3 mismatches. **c**, Snapshots showing CHH, CHG and CG methylation at MetGene186 in meiocytes from wild type (WT) and two independent CRSIPR lines carrying deletions (shown in Supplementary Fig. 2a) of HyperTE224. MetGene186 is underlined in red, as in Fig. 1b. **d**, Bar plots displaying DNA methylation levels at MetGene24 (*AT1G15520*) and 86 (*AT1G56410*) in meiocytes from WT, two independent CRISPR lines carrying HyperTE315 (*AT2TE72995*) deletions (shown in Supplementary Fig. 2d), or the *rdr2* or *drm* mutant.

To test the causal relationship between HyperTE-associated siRNAs and the methylation at MetGenes, we created two independent CRISPR/Cas9 lines with deletions of a HyperTE (HyperTE224, *AT2TE15980*; Fig. 2b and Supplementary Fig. 2a). We observed abolishment of methylation at the predicted target MetGene (MetGene186, *AT3G04230*; Fig. 2b and Supplementary Table 3) in meiocytes isolated from both deletion lines (Fig. 2c), whereas overall methylation did not change (Supplementary Fig. 2b). We applied this experimental strategy to another HyperTE (HyperTE315, *AT2TE72995*), siRNAs from which are predicted to target two MetGenes (MetGene24, *AT1G15520;* and MetGene86, *AT1G56410*) (Supplementary Fig. 2c and Supplementary Table 3). Consistently, we observed complete loss of methylation at both target MetGenes in meiocytes of the HyperTE315-deletion lines (Fig. 2d and Supplementary Fig. 2b,d). These results unambiguously demonstrate that methylation at MetGenes is induced *in trans* by siRNAs derived from HyperTEs.

### Meiocyte siRNA profiles closely resemble those of the tapetal nurse cells

As RdDM is self-reinforcing, 24-nt siRNAs bear no mismatches with their targets, because regardless of the initial guiding siRNA, Pol V-mediated DNA methylation attracts the Pol IV pathway *in situ* to generate perfectly matching siRNAs^5,6^. The observation that very few perfectly matching siRNAs are associated with MetGenes (Figs. 1b,c and 2a) suggests that the Pol IV and Pol V pathways are decoupled in meiocytes, with MetGene-inducing siRNAs generated from HyperTEs in other cells.

Male meiocytes are completely enclosed by a layer of tapetal nurse cells^19^ (Fig. 1a). The meiocytes and tapetal cells are extensively connected via cytoplasmic channels called plasmodesmata^26,27^, which are widely accepted as the most likely route for intercellular siRNA movement in plants^28–30^. Furthermore, 24-nt siRNAs have been proposed to accumulate in maize tapetum during early meiosis^31^, exactly the stage when plasmodesmata connect meiocytes and tapetal cells. Therefore, siRNAs that induce methylation at MetGenes may be generated by HyperTEs in the tapetum and transported into meiocytes. To test this hypothesis, we developed a method to isolate tapetal cells. We generated *Arabidopsis* transgenic plants carrying GFP driven by a tapetum-specific promoter (*pA9*, simplified as *pTP*; Fig. 3a)^32–36^ and isolated GFP-positive tapetal cells via fluorescence-activated cell sorting (FACS; Supplementary Fig. 3a). The isolated tapetal cells are of high purity, evaluated through fluorescence microscopy (>95% purity) and RNA sequencing, which shows enrichment of tapetum-specific genes and depletion of somatic and meiotic genes (Supplementary Table 4).

**Fig. 3:**
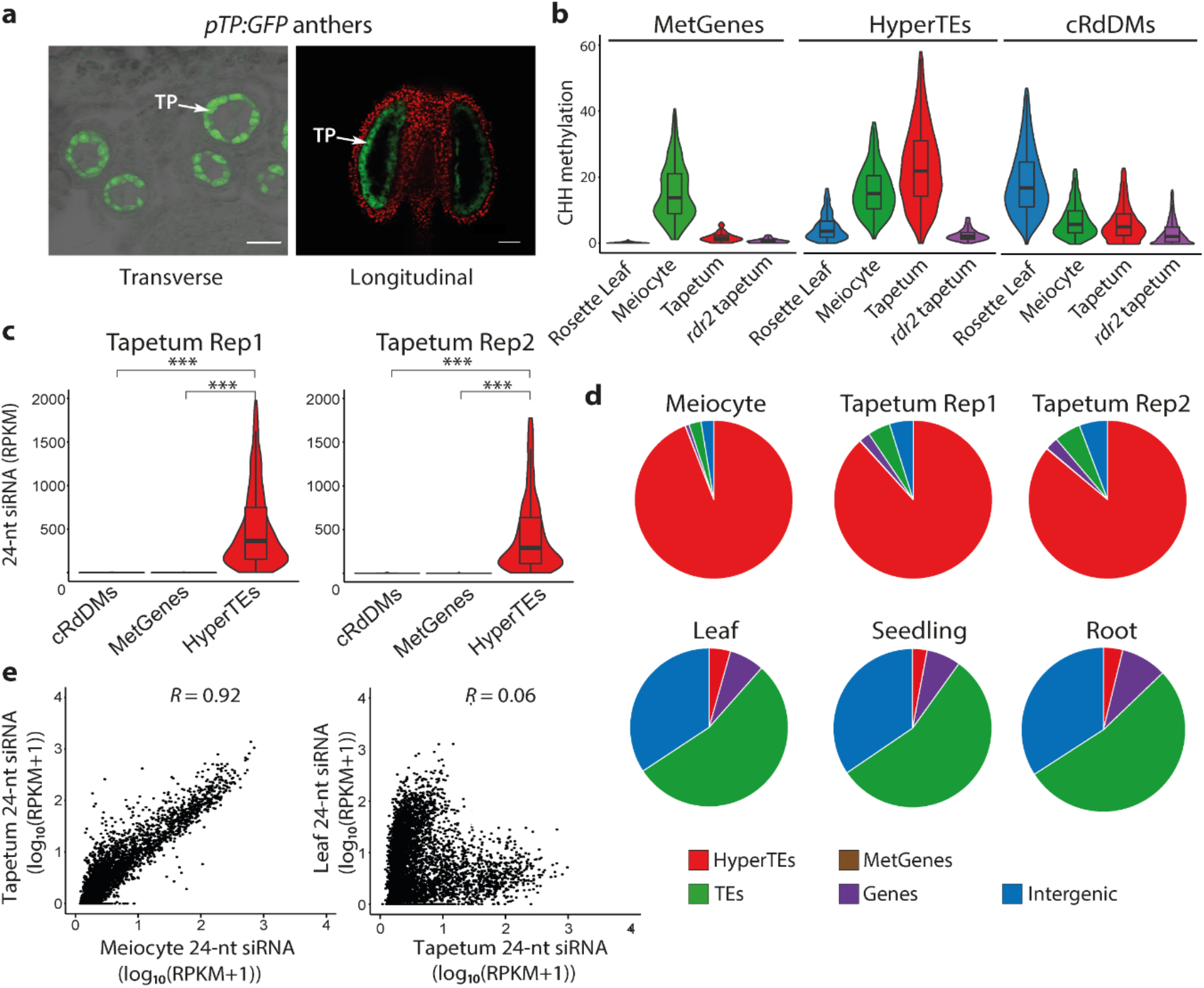
Tapetal nurse cells have a similar 24-nt siRNA profile to meiocytes. **a**, Transverse (differential interference contrast, DIC) or longitudinal (confocal) image of anthers with the tapetum (TP) labelled by *pTP::GFP*. Green, GFP fluorescence; red, auto-fluorescence. Scale bars: 25 μm. **b**, Violin plots showing CHH methylation at MetGenes, HyperTEs or cRdDMs in the WT and *rdr2* mutant tapetum, in comparison to WT leaf and meiocyte. **c**, Violin plots illustrating 24-nt siRNA levels at cRdDMs, MetGenes and HyperTEs in the tapetum (two biological replicates), as in Fig. 1c. *** *P* < 2.2e- 16, Kolmogorov-Smirnov test. **d**, Pie charts depicting the proportions of 24-nt siRNAs associated with different genomic annotations in the tapetum (two biological replicates), compared to the meiocyte and somatic tissues (leaf, seedling and root). **e**, Scatter plots showing the correlation between 24-nt siRNA abundance in the tapetum and meiocyte at meiocyte siRNA clusters (left; Pearson’s *R*=0.92, n=3,712), and between that in the rosette leaf and tapetum over tapetum siRNA clusters (right; Pearson’s *R*=0.06, n=5,841).

DNA methylation analysis of tapetal cells revealed that HyperTEs are hypermethylated in the tapetum in comparison to somatic tissues, at levels even higher than in meiocytes (Figs. 1b and 3b). As in meiocytes (Fig. 1b,d), HyperTE methylation in the tapetum requires RDR2 (Figs. 1b and 3b). Consistently, tapetal cells show substantial enrichment of 24-nt siRNAs at HyperTEs (Fig. 3c,d and Supplementary Fig. 3b), similar to meiocytes (Fig. 1c). As in meiocytes (Fig. 1c), siRNA levels at canonical RdDM loci are relatively low but similar in absolute terms to somatic cells (Figs. 1e and 3c) and DNA methylation at these loci is low (Figs. 1b and 3b). Genome-wide, 24-nt siRNAs in identified siRNA clusters strongly correlate between tapetal cells and meiocytes (Pearson’s *R* = 0.92; in contrast, *R* = 0.06 between tapetum and leaf; Fig. 3e). As in meiocytes (Fig. 1c), few siRNAs match MetGenes perfectly in tapetal cells (Fig. 3c). The strong resemblance between siRNA profiles suggests *en masse* siRNA movement from tapetum to meiocytes, which is consistent with knowledge of intercellular siRNA transport through plasmodesmata^37^. Importantly, in contrast to meiocytes, MetGenes are not methylated in tapetal cells (Figs. 1b and 3b). This is consistent with tapetum having an active Pol IV pathway, because RdDM-mediated methylation should not exist in the absence of perfectly matching siRNAs in such cells. Our results also imply that, unlike in meiocytes, the tapetal Pol V pathway is unable to induce DNA methylation with mismatched siRNAs.

### Nurse cell-derived siRNAs (niRNAs) drive meiocyte methylation reprogramming

To test whether tapetum-derived siRNAs work *in trans* to induce methylation in meiocytes, we created a genetic mosaic system with 24-nt siRNA biogenesis confined to the tapetum. We expressed RDR2 (fused with a small Flag tag) in the *rdr2* null mutant background using the tapetum-specific *pA9* promoter (simplified as *pTP::RDR2 rdr2*; Fig. 4a). Immunolocalization with anti-Flag antibodies in anther cross-sections confirmed the specific induction of RDR2 in the tapetum in two independent mosaic lines (Fig. 4a). In the meiocytes isolated from both lines, methylation at MetGenes and HyperTEs was increased to levels comparable with wild type (Fig. 4b,c and Supplementary Fig. 4a,b). As a control, we generated a *pTP::POL-V pol-v* mosaic line. As Pol V mediates DNA methylation rather than siRNA biogenesis, it has to work *in cis*. Consistently, meiocytes from two independent lines show no methylation at MetGenes (Fig. 4b,c; Supplementary Fig. 4c shows Pol V protein is correctly expressed). These results demonstrate that activity of the Pol IV pathway in the tapetum is sufficient to methylate MetGenes and HyperTEs in meiocytes.

**Fig. 4:**
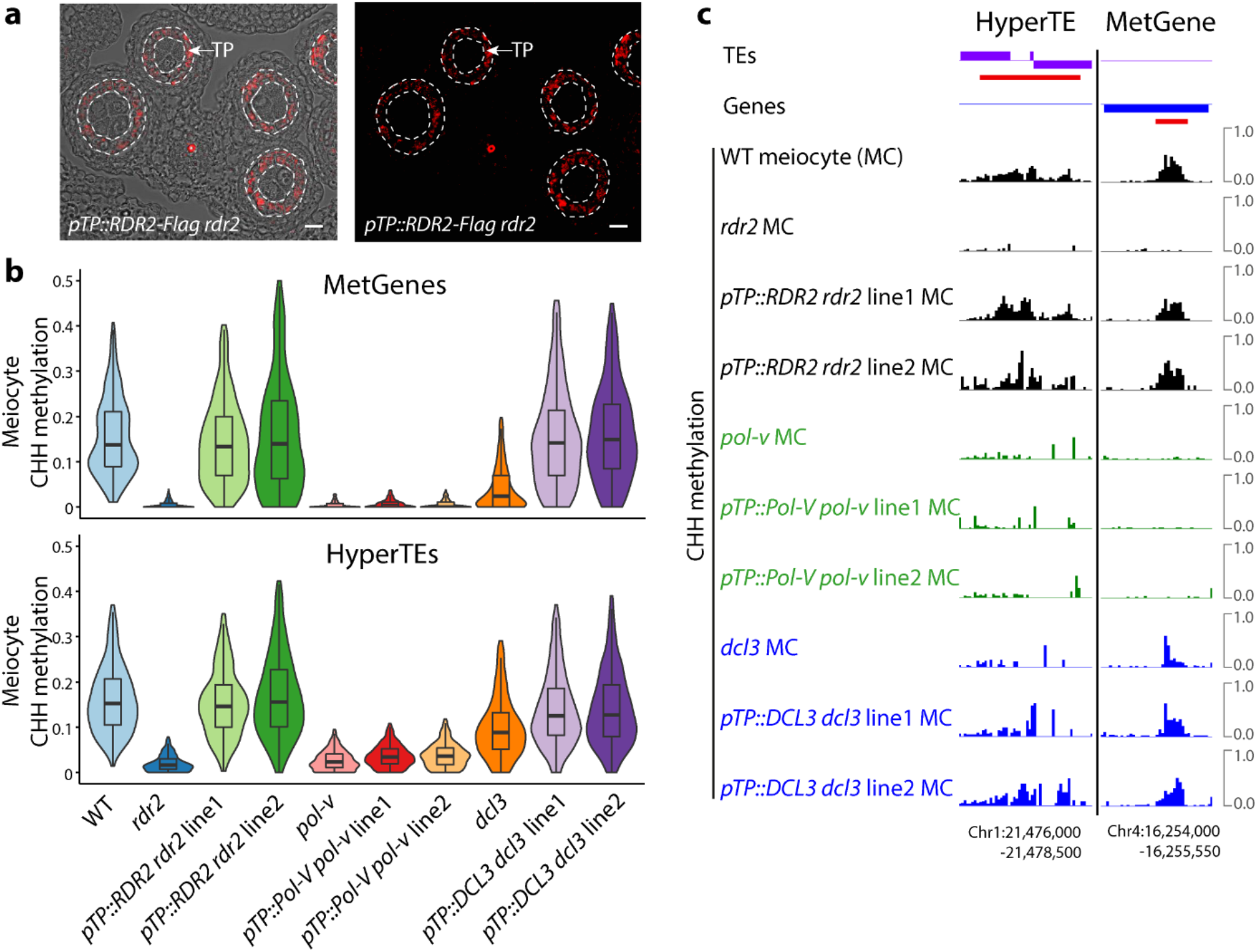
siRNAs that drive methylation in meiocytes are produced by tapetal nurse cells. **a**, Immunostaining of *pTP::RDR2-Flag rdr2* anther cross-sections using anti-Flag antibody. Signal (red) is observed specifically in the tapetum (TP, outlined in white) and imaged in the transmitted light (DIC, left) or dark field (right). Scale bars: 10 μm. **b**, Violin plots showing CHH methylation at MetGenes (upper panel) and HyperTEs (lower panel) in the meiocytes from WT, RdDM mutants and their corresponding genetic mosaic lines (two independent transgenic lines are shown for each). **c**, Snapshots of CHH methylation at a HyperTE and a MetGene (underlined in red) in the meiocyte (MC) from different genotypes (the same MC genotypes are depicted in **b**).

RDR2 produces double-stranded RNA (dsRNA) precursors, which are converted to 24-nt siRNAs by DCL3^5^. Because the dsRNA precursors have been shown to move between cells^28,29^, we sought to establish whether these molecules or siRNAs move from the tapetum to meiocytes. We therefore generated two independent DCL3 mosaic lines (*pTP::DCL3 dcl3*; Supplementary Fig. 4c shows DCL3 is correctly expressed). Meiocytes from both lines show fully wild-type methylation at MetGenes and HyperTEs, whereas *dcl3* mutants have significantly reduced methylation at both sets of loci in meiocytes (both *P* < 2.2e-16, Kolmogorov-Smirnov test; Fig. 4b,c). This result indicates that 24-nt siRNAs are the non-cell-autonomous signal that originates in the tapetum and drives methylation at MetGenes (and HyperTEs) in meiocytes. Therefore, we will refer to these as <u>n</u>urse-cell-der<u>i</u>ved siRNAs (niRNAs) hereafter.

### Nurse-cell-specific CLSY3 is required for niRNA transcription from HyperTEs

We next investigated how the distinctive niRNA profile in the tapetum is generated (Fig. 3c,d). Four putative chromatin remodelers, CLASSY1-4 (CLSYs), recruit Pol IV to produce siRNAs at largely discrete sets of loci^38^. Therefore, we evaluated the relationship of the *clsy1*/*2*/*3*/*4*-dependent siRNA clusters identified in floral buds^38^ with HyperTEs. We found a substantial proportion of HyperTEs (66%; 524 out of 797) overlap *clsy3*-dependent siRNA clusters (804 loci), whereas only 1% to 8% of HyperTEs overlap *clsy1*-, *2*- or *4*-dependent clusters (Supplementary Fig. 5a). Consistently, in both tapetal cells and meiocytes, relatively few 24-nt siRNAs are associated with *clsy1,2*- or *clsy4*-dependent clusters, whereas *clsy3*-dependent clusters are significantly enriched in siRNAs (all *P* < 2.2e-16, Kolmogorov-Smirnov test; Fig. 5a).

**Fig. 5:**
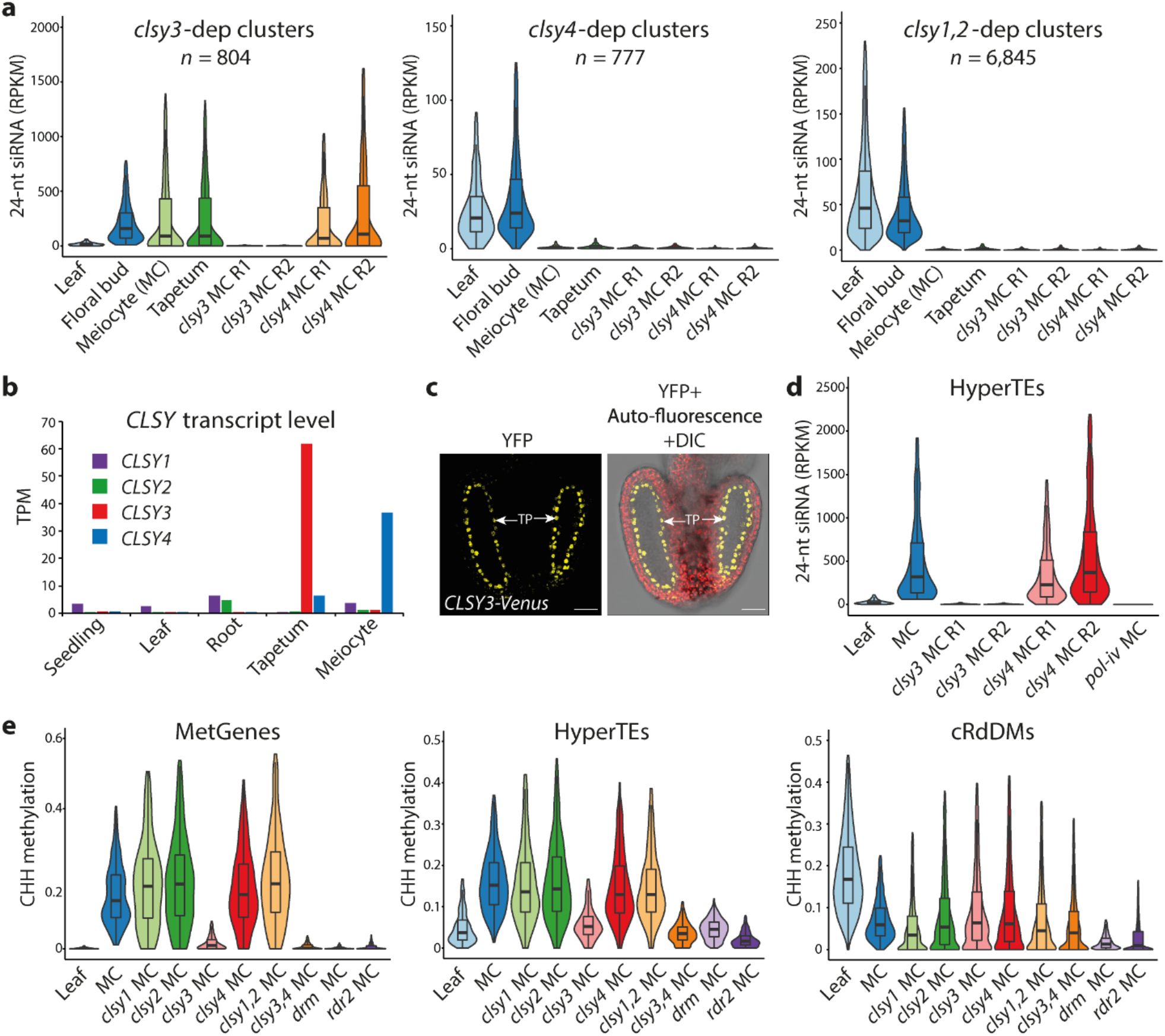
Tapetum-specific CLSY3 is responsible for the production of niRNAs. **a**, Violin plots depicting 24-nt siRNA abundance at published *clsy3*-, *clsy4*-, *clsy1*,*2*-dependent (dep) clusters in the leaf, floral bud, meiocyte (MC) and tapetum. R, biological replicate. **b**, Abundance of *CLSY* transcripts in the seedling, rosette leaf, root, tapetum and meiocyte. TPM, transcripts per million. **c**, Confocal images showing the specific localization of Venus-tagged CLSY3 in the tapetum (TP) at anther stage 6, when meiosis is ongoing^87^. Scale bars: 25 μm. **d**, Violin plots illustrating 24-nt siRNA abundance at HyperTEs in the leaf and meiocyte (MC) from WT, *clsy3*, *clsy4*, and *pol-iv* mutants. **e**, Violin plots showing CHH methylation at MetGenes, HyperTEs and cRdDMs in the leaf and the meiocyte (MC) from WT and different mutant plants.

We next examined the transcription of *CLSY*s in the tapetum, meiocytes and somatic tissues. We found that *CLSY1* and *2* are the predominant homologs in somatic tissues, whereas *CLSY3* is by far the most highly expressed *CLSY* in the tapetum (72.2 TPM; Fig. 5b). To validate this result and examine the expression pattern of CLSY3, we generated a *CLSY3-Venus* fusion line (*pCLSY3::Venus-CLSY3*). Confocal microscopy shows CLSY3 is undetectable in seedlings and leaves but enriched in carpels and anthers (Fig. 5c and Supplementary Fig. 5b). In the anther, CLSY3 is specifically expressed in tapetal cells and absent from all other cell types, including meiocytes (Fig. 5c). Based on these results, CLSY3 is likely responsible for the distinctive siRNA profile in tapetal and meiocyte cells.

To test this hypothesis, we isolated meiocytes from the *clsy3* mutant and performed siRNA sequencing. We also examined *clsy4* meiocytes, because *CLSY3* and *4* were reported to be partially redundant^38^ and *CLSY4* transcripts are detected in the tapetum (6.6 TPM; Fig. 5b). We found that 24-nt siRNAs at HyperTEs are almost completely lost in *clsy3* mutant meiocytes, similarly to *pol iv* mutant meiocytes, whereas *clsy4* meiocytes are unaffected (Fig. 5d). Consistently, we found greatly reduced methylation at HyperTEs and MetGenes in *clsy3* meiocytes, whereas *clsy1*, *2* or *4* mutant meiocytes and *clsy1;2* double mutant meiocytes have wild-type levels of methylation (Fig. 5e). These results establish that CLSY3 is required for niRNA production and the consequent methylation of HyperTEs and MetGenes. Given that CLSY3 is absent from meiocytes and only expressed in the tapetum (Fig. 5c), these results further demonstrate that siRNA biogenesis is suppressed in meiocytes and meiocyte siRNAs are derived from the tapetum.

### Tapetal niRNAs drive methylation reprogramming in the entire male germline

As the methylation at MetGenes persists throughout male germline development to gametogenesis^11^ (Fig. 1b), we wondered whether tapetal niRNAs can drive methylation throughout the germline. To test this, we analyzed DNA methylation in sperm isolated from the *clsy3* mutant. CHH methylation at MetGenes and HyperTEs is reduced in *clsy3* mutant sperm to levels resembling *drm* mutant sperm (Fig. 6a,b). In contrast, methylation at canonical RdDM loci is not affected by the *clsy3* mutation (Fig. 6a,b and Supplementary Fig. 6).

**Fig. 6:**
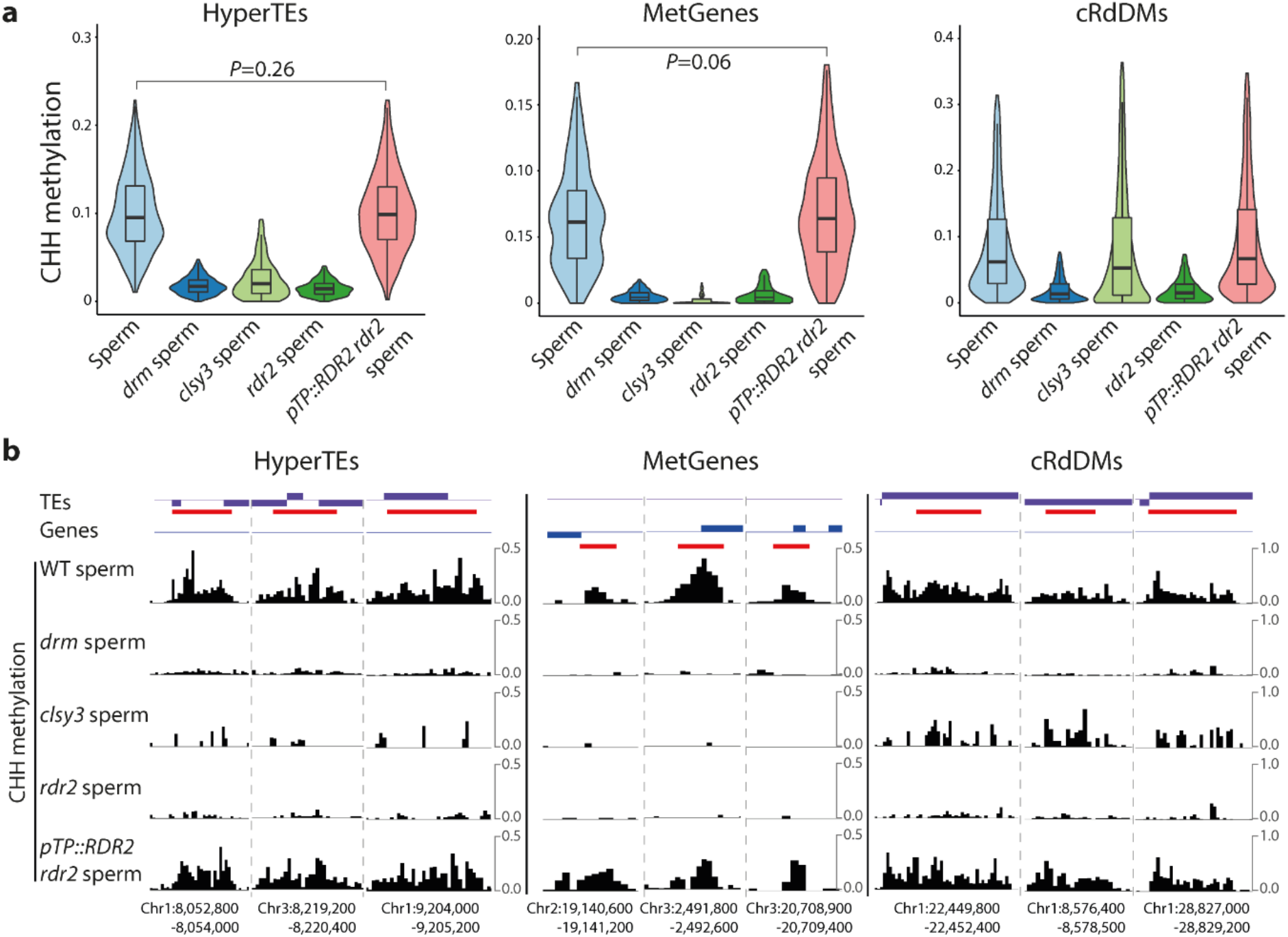
Tapetal niRNAs determine the sperm DNA methylome. **a**, Violin plots showing CHH methylation at HyperTEs, MetGenes, and cRdDMs in the sperm from WT and different genotypes. *P* values, Kolmogorov-Smirnov test. **b**, Snapshots of CHH methylation at example HyperTEs, MetGenes, and cRdDMs (all underlined in red) in the sperm from different genotypes.

To confirm the competence of tapetal niRNAs to shape sperm methylation, we examined sperm from the *pTP::RDR2 rdr2* mosaic line. DNA methylation at MetGenes and HyperTEs is restored to wild-type levels in *pTP::RDR2 rdr2* sperm (Fig. 6a,b and Supplementary Fig. 6). Furthermore, methylation at canonical RdDM loci is also restored (Fig. 6a,b and Supplementary Fig. 6), in line with soma-like absolute levels of 24-nt siRNA at canonical RdDM loci in the tapetum (Fig. 1e). These results demonstrate that tapetal niRNAs drive DNA methylation reprogramming in the male germline all the way to the gametes.

### Tapetal niRNAs silence transposons in the germline

As tapetal niRNAs induce methylation at TEs in the male germline (Figs. 4b, 5d and 6a), we investigated whether niRNAs serve to silence germline TEs in addition to their demonstrated gene regulatory function^11^. To address this question, we performed RNA sequencing of tapetal cells and pollen from wild type and the *drm* mutant (Fig. 1a). Analysis of these data in combination with analogous data from meiocytes and leaves^11^ identified retrotransposons from a Gypsy family (*GP1*) that are specifically activated in RdDM-defective germline cells (Fig. 7a). *GP1* transcription is not activated in *drm* mutant leaves, but strongly activated in the tapetum and pollen (and slightly in meiocytes; Fig. 7a). We observed loss of methylation in RdDM (*drm* and *rdr2*) mutants at the LTRs in the tapetum and meiocytes (Supplementary Fig. 7a), indicating that *GP1* is directly suppressed by RdDM.

**Fig. 7:**
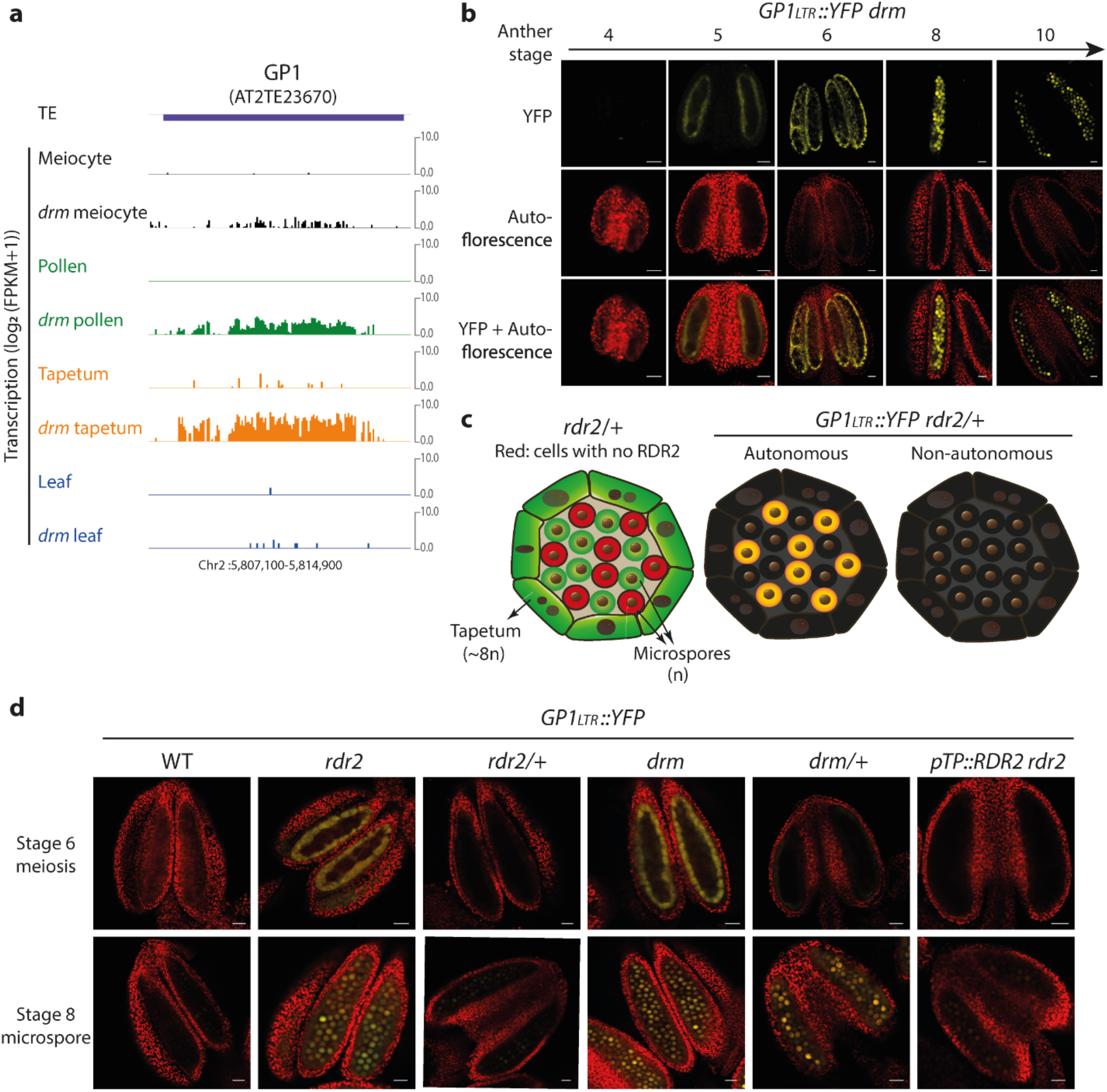
Tapetal niRNAs silence the *GP1* transposon in the germline. **a**, Snapshots of transcription at *GP1* in the meiocyte, pollen, tapetum and leaf from WT and the *drm* mutant. **b**, Confocal images of *GP1_LTR_::YFP drm* anthers during development. The time axis shows anther stages^87^ from early to late. TP, tapetum; MSP, microspore. Yellow, YFP; red, auto-fluorescence. Scale bars: 20 μm. **c**, Cartoons of anther locules depicting the experiment in **d** (left, cellular RDR2 functionality in *rdr2*/+ heterozygous mutant) and the expected outcome if *GP1*-silencing siRNA is cell autonomous (middle) or non-autonomous (right). **d**, Anther confocal images of *GP1_LTR_::YFP* in different genetic backgrounds, taken at anther stage 6 (upper panels) and 8 (lower panels). As in **b**, YFP and auto-fluorescence in yellow and red, respectively. Scale bars: 25 μm.

To further examine *GP1* activity during germline development, we generated a YFP reporter line driven by *GP1*’s LTR sequence (*GP1_LTR_::YFP*) and crossed it to *rdr2* and *drm* mutants. Consistent with the RNA-seq data, YFP signal was undetectable in wild type and specifically observed in the anthers of *rdr2* and *drm* mutants (Fig. 7b,c,d). In both mutants, *GP1* activity can be first observed in the tapetum at a stage prior to meiosis (Fig. 7b). Subsequently, YFP signal is strongest in the tapetum at the onset of meiosis, and in microspores (and to a lesser extent pollen) after meiosis (Fig. 7b,c,d). These observations confirm that *GP1* is specifically activated in the germline and tapetum when RdDM is compromised.

To confirm that tapetum is the origin of *GP1* silencing siRNAs, we first took advantage of the fact that microspores are haploid meiotic products in which genetic segregation has occurred (Fig. 1a). In a mutant carrying a recessive heterozygous *rdr2* mutation, the diploid tapetal cells (of *rdr2*/+ genotype) are able to produce siRNAs, whereas half of the haploid microspores (the ones of *rdr2* genotype) are not (Fig. 7c). We did not detect *GP1* activity in any anther tissue, including the tapetum and microspores, of *GP1_LTR_::YFP rdr2/+* plants (Fig. 7d), confirming that *GP1*-silencing siRNAs are produced in diploid somatic cells. In contrast, half (50.7%, n = 1391) of the microspores from *GP1_LTR_::YFP drm/+* plants exhibit YFP signal (Fig. 7d), consistent with the *cis* action of DRM methyltransferases.

To validate the silencing of *GP1* by tapetal niRNAs, we crossed the *GP1* YFP reporter into the *pTP::RDR2 rdr2* genetic background that limits siRNA production to the tapetum. No *GP1* activity was observed in *GP1_LTR_::YFP pTP::RDR2 rdr2* anthers (Fig. 7d), demonstrating the competence of tapetal niRNAs to silence *GP1* in the male germline. Importantly, *GP1* is a canonical RdDM locus (Supplementary Fig. 7a,b), further demonstrating that tapetal niRNA production and germline activity are not confined to HyperTEs.

## Discussion

### Meiocytes can methylate genes that imperfectly match TE-derived siRNA

How the TE-focused RdDM pathway can target specific genes has been a major mystery regarding DNA methylation reprogramming in the male germline. Our results demonstrate that male meiocytes can induce DNA methylation of genes that imperfectly match siRNAs. This off-targeting likely requires large quantities of siRNAs, because the methylated MetGenes have sequences that correspond to HyperTEs, which account for most of the siRNAs in meiocytes (Fig. 1c,f). However, it is noteworthy that HyperTE siRNA accumulates to similar levels in the tapetum (Fig. 3c,d), yet cannot target methylation of MetGenes (Figs. 3b and 8). This difference could have two explanations: the Pol V branch of RdDM might be hypersensitively configured in meiocytes, or meiocyte chromatin might permit Pol V targeting by imperfectly matching siRNAs. These explanations are not mutually exclusive, and are consistent with the observation that tethering Pol V pathway components is sufficient to induce methylation even in the absence of siRNAs^39^. Investigating the meiocyte-specific features of Pol V functionality will be an important focus of future studies.

The finding that genic methylation in the male germline is targeted by TE-produced siRNAs suggests that the germline gene regulatory functions of RdDM have evolved from the pathway’s main TE silencing activity. The finding that Gypsy *GP1* retrotransposons are suppressed by germline RdDM supports this idea. Loss of RdDM does not activate *GP1* in somatic tissues (Fig. 7a), whereas DNA methylation of the LTRs requires RdDM in the soma, tapetum and germline (Supplementary Fig. 7a). This suggests that *GP1* specifically targets expression in reproductive cells, possibly by exploiting transcription factors specific to these cell types. This may be advantageous for the transposon, as expression in somatic tissues would not contribute to TE inheritance but could trigger silencing via systemic RdDM^40–44^. The aggressive configuration of the Pol V pathway in meiocytes may have evolved to counteract such TEs.

### Tapetal niRNAs drive germline methylation reprogramming

Our results indicate that male meiocytes combine a highly sensitive Pol V pathway with suppression of the Pol IV pathway (Fig. 8). Although our data cannot rule out that some siRNA production occurs in meiocytes, the observation that meiocyte siRNAs are strongly dependent on the tapetum-specific Pol IV co-factor CLSY3 (Fig. 5d,e) indicates that the vast majority of meiocyte 24-nt siRNAs are derived from the tapetum (Fig. 8). The Pol IV pathway may be suppressed in meiocytes because siRNA production, which involves transcription, carries the risk of TE activation. Biogenesis of siRNAs might be carried out more safely by the short-lived tapetum.

**Fig. 8:**
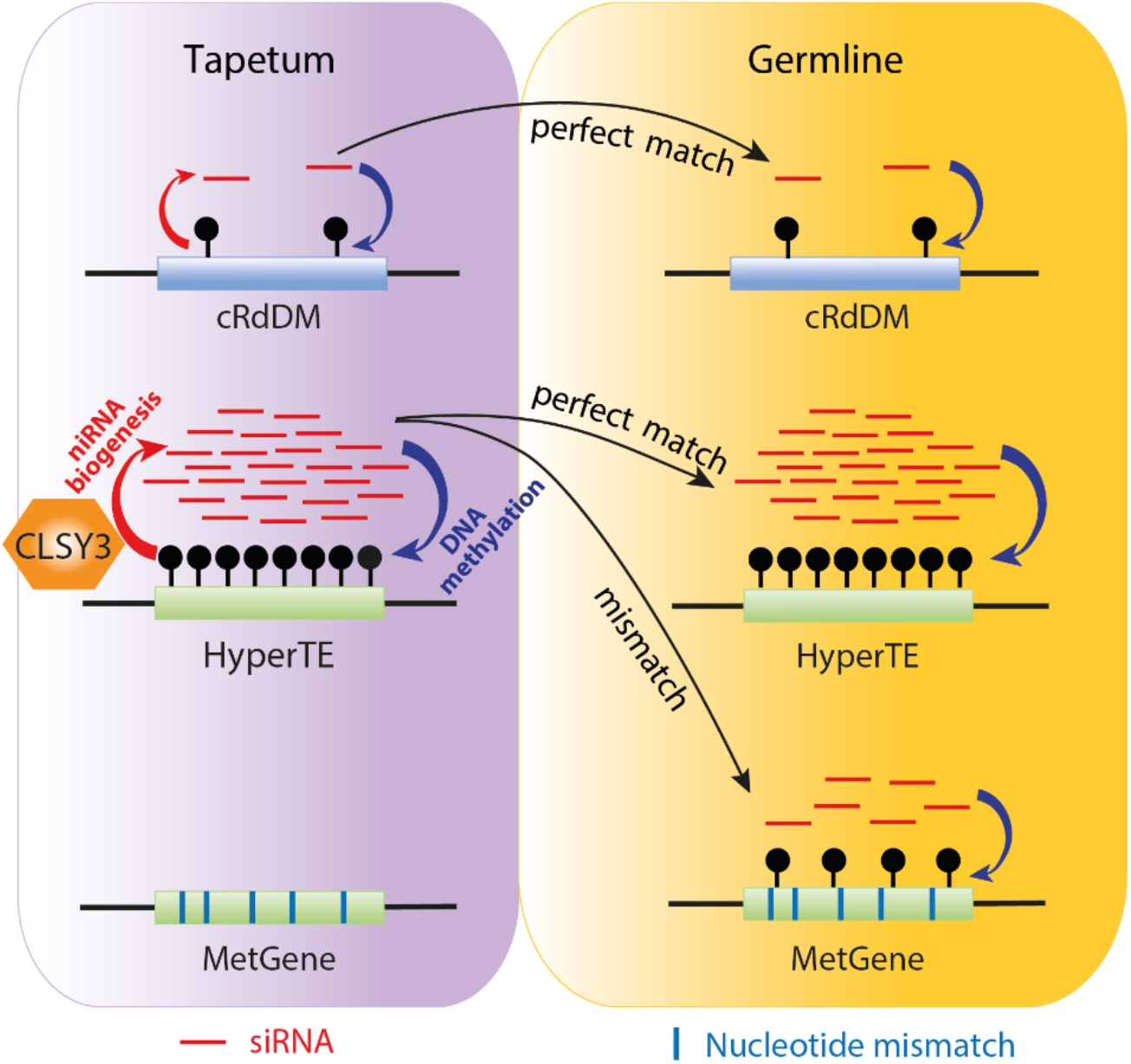
Cartoon depicting the establishment of germline methylation by tapetal niRNAs. Tapetal niRNAs are predominantly transcribed from HyperTEs, enabled by the specific expression of the Pol IV recruiter CLSY3. These HyperTE-derived niRNAs act non-cell-autonomously in the male germline to drive hypermethylation at perfectly matched HyperTEs, and induce methylation at MetGenes with similar sequences. Tapetal niRNAs are capable of driving methylation in the germline from meiocytes to the end product, sperm, even at canonical RdDM loci (cRdDMs) that associate with fewer niRNAs than HyperTEs do. MetGene methylation is germline-specific, suggesting that a specialized Pol V pathway and/or chromatin state permits methylation via mismatched niRNAs. Accordingly, MetGenes in tapetal cells are not methylated, despite the high abundance of HyperTE-derived niRNAs.

Remarkably, tapetal niRNAs can induce DNA methylation not only in meiocytes, but throughout the male germline (Fig. 6a). Our data show that mutation of *CLSY3* abolishes MetGene methylation in sperm (Fig. 6a), and the *pTP::RDR2 rdr2* mosaic line with siRNA biogenesis confined to the tapetum restores not only sperm MetGene methylation, but also methylation at canonical RdDM loci (Fig. 6a) and *GP1* silencing (Fig. 7d). As direct siRNA movement between pollen and tapetum is hampered by the pollen wall^20^, niRNAs most likely influence sperm via inheritance from meiocytes. Previously, siRNAs from the pollen vegetative cell were shown to reinforce TE methylation in sperm^23,45^. The sperm methylation landscape is therefore likely determined by two waves of exogenous siRNAs, first from the tapetum and another from the vegetative cell. The tapetum plays a central role in this reprogramming process, as tapetal niRNAs are required to establish MetGene methylation and are competent to drive the full spectrum of sperm RdDM (Figs. 6a and 8).

Tapetal cells are descendants of the same progenitor cell as meiocytes and specialize in transporting biological materials to meiocytes^19^. Thus, they are ideal for genome surveillance of TEs and the production of siRNAs. In meiocytes, the Pol V pathway is tuned aggressively, so that TEs that produce siRNAs in the tapetum and ones with similar sequences are targeted. The hypersensitivity of meiocyte RdDM also causes methylation of genes bearing similar sequences (Fig. 8), which potentiates transcriptional regulation and hence the selection of beneficial functions during evolution^11^.

### Tapetal niRNAs closely resemble phasiRNAs

Our results indicate that niRNAs accumulate in the tapetum at the pachytene stage of meiosis. Other plant species accumulate 24-nt phased siRNAs (phasiRNAs) in the tapetum at a similar developmental stage^31,46–49^. The phasiRNA clusters of monocot species such as maize and rice differ in their biogenesis from niRNAs, as they are transcribed by Pol II and are processed by a monocot-specific family of DCL nucleases (DCL5)^48,50–52^. Nonetheless, like *Arabidopsis* niRNAs, phasiRNAs are transcribed from hundreds of loci and are highly abundant, accounting for 64% of all 24-nt siRNAs in maize anthers^31,50,52,53^. Comparable to the importance of niRNAs for meiotic progression^11^, phasiRNAs are important for fertility in maize and rice^51,53,54^. How phasiRNAs function remains unknown due to the lack of perfectly matching genomic targets^55,56^. Whether phasiRNAs function primarily in the tapetum or are transported into meiocytes is also unclear^31,49^. Our discovery that *Arabidopsis* niRNAs target mismatched genes in meiocytes and regulate TEs in the male germline opens new possibilities for understanding phasiRNA function.

### Evolutionary convergence of the plant niRNA and animal piRNA pathways

The piRNA pathway, which operates specifically in germ cells, forms a conserved RNA interference mechanism for silencing TEs during animal sexual reproduction^57^. Plants have a variety of siRNA pathways that are functionally important for plant development and reproduction^58–65^. However, plants lack piRNAs, as the Piwi clade of Argonaute nucleases is restricted to metazoans^66^. Whether plants have a functionally equivalent germline RNAi pathway has been unclear. 21-nt siRNAs, typically involved in post-transcriptional silencing, have been implicated in the movement from the pollen vegetative cell to sperm and the silencing of sperm TEs^45,67^. However, the ability of endogenous vegetative cell siRNAs to mediate silencing in sperm has not yet been demonstrated, the nature and silencing mechanism of the non-cell autonomous signal have remained controversial, and no RNAi genes have been specifically implicated^68–71^. The general plant RdDM pathway has been suggested to share deep homology with piRNA pathways^66^, but RdDM functions throughout plant tissues, unlike piRNA pathways that operate only in germ cells. Whether plants have specifically configured germline RNAi pathways has thus remained unknown.

Our study demonstrates the presence of a specialized niRNA pathway in plant germ cells. The niRNA pathway bears strong similarities to metazoan piRNAs in terms of biogenesis and function. Like piRNAs, niRNAs are important for fertility, specifically enriched in reproductive cells (Fig. 3d), and capable of silencing TEs (Fig. 7d). niRNAs are produced in tapetal nurse cells and transported into meiocytes, like *Drosophila* piRNAs that are transported from nurse cells into oocytes^12^. The ability of niRNAs from TE clusters to regulate mismatched genes *in trans* is similar to the described silencing of male genes by a piRNA transcribed from repeats on the silkworm female sex chromosome^72^. Intriguingly, mammalian piRNAs are enriched in pachytene spermatocytes (the same developmental stage as niRNAs) and lack perfectly-matched mRNA targets^57,73^. Mouse pachytene piRNAs have been shown to target and post-transcriptionally regulate mismatched genes^74,75^, as have piRNAs in *Drosophila* and *C. elegans*^57,74^. The recurrence of broad targeting competence predicts a general ability of niRNAs and piRNAs to regulate genes as well as TEs. Overall, the many similarities between niRNAs and piRNAs indicate that gametogenesis in plants and animals requires specialized small RNA pathways to control TEs and preserve genome integrity, and these pathways evolve to regulate gene activity and fertility.

## Methods

### Plant materials

All plants used in this study (*Arabidopsis thaliana* Col-0) were grown under 16h light/ 8h dark in a growth chamber (21°C, 70% humidity). The following published lines were used in this study: *clsy1-7*, *clsy2-1*, *clsy3-1* and *clsy4-1*^38^, *rdr2-1*^76^, *nrpe1-11* (*pol-v*)^77^, *dcl3-1*^78^*, drm1-2 drm2-2* (*drm*)^79^. *clsy1,2* double and *clsy3,4* double mutants were created by crossing the corresponding single mutants. Transgenic plants were obtained by agrobacterium-mediated floral dip. The *pA9::GFP* vector was transformed into wild-type (WT) plants: the T2 and T3 transgenic plants were used for experiments, and the T1 transgenics were also crossed with the *rdr2* mutant to obtain *pA9::GFP rdr2*. *GP1_LTR_::YFP* was transformed into WT: the resulting T2 plants were used in experiments and crossed with the *rdr2* mutant, the *drm* mutant and the *pA9::RDR2 rdr2*. Vectors *pA9::RDR2-Flag*, *pA9::Pol-V-3×Flag*, and *pA9::DCL3-3×Flag* were transformed into, respectively, the *rdr2*, *pol-v*, *and dcl3* mutant backgrounds; the *pCLSY3::Venus-CLSY3* vector was transformed into the *clsy3* mutant. For these, two independent T1 lines were selected for each, and the T2 and T3 generations were used for experiments.

### Isolation of male meiocytes, tapetal cells and sperm

Male meiocytes (prophase I) and sperm nuclei were isolated as described previously^11^.

To obtain tapetal cells, flower buds were harvested from the *pA9::GFP* and the *pA9::GFP rdr2* lines and protoplasted as previously described^80^, except that the enzymatic digestion of cell wall was performed for only 1h. Prepared protoplasts were gently suspended in a buffer containing 0.2 M mannitol, 12.5 mM KCl, 67.5 mM CaCl_2_, 11 mM MES (pH 5.7), 0.05% BSA (w/v) and 50 mM NaCl, and sorted on a BD FACSMelody cell sorter (Beckton Dickinson). The sorting was performed using a 70 μm ceramic nozzle with 1× PBS running at a constant pressure of 20 psi. GFP-positive tapetal protoplasts (refer to Supplementary Fig. 3a for the gate setting) were collected, examined under a fluorescence DM6000 microscope (Zeiss) for purity, and stored immediately at −80°C.

### Plasmid construction

To construct the *pA9::GFP* vector, *A9* promoter (*pA9*)^33^ was cloned using primers PHG1 and PHG2 and inserted into the destination vector *pDMC107-NTF*^81^ using the Gateway system (Invitrogen, cat. no. 11791-020).

For the *pA9::RDR2*-*Flag* construct, *pA9* and *RDR2* (the ORF without the stop codon) were amplified using primer pairs PHG001-PBA008 and PBA009-PBA010 respectively. The PCR products were then combined to amplify *pA9::RDR2* using primers PHG001 and PBA010. Finally, *pA9::RDR2* was introduced into the destination vector *pGWB10*^82^, which contains a *Flag* tag, using Gateway.

For the *pA9::Pol-V-3×Flag* plasmid, *pA9* was cloned using primers PJL101 and PJL102, and the ORF of *Pol-V* (*NRPE1*, without the stop codon) was cloned with primer PJL104 and PJL105 (containing 3*×Flag* sequences). These PCR products were then cloned into the *pCAMBIA1301* vector (linearized by *PstI* and *NcoI* restriction enzymes, New England BioLabs, cat. no. R3140 and cat. no. R3193) using the In-fusion (Takara, cat. no. 638909) method. Similarly, the *pA9::DCL3-3×Flag* construct was made using primers PJL101, PJL103, PJL106 and PJL107.

For the *pCLSY3::Venus-CLSY3* vector, primer pairs JW568-JW569 and JW576-JW577 were used to amplify *pCLSY3* and *CLSY3*, respectively. Amplified products were cloned into the donor vector *pK7m34GW*^83^, and subsequently the destination vector *pGWB13-Bar*^11^ using the MultiSite Gateway method (Invitrogen, cat. no. 11791-020).

For the *GP1_LTR_::YFP* construct, the 5’LTR sequence of *GP1* (*AT2TE23670*) was cloned into the donor vector *pDONR207* (Invitrogen) using primers PHG121 and PHG124. *GP1_LTR_* was then cloned into the *YFP*-containing destination vector *pGWB40*^82^ using the MultiSite Gateway (Invitrogen, cat. no. 11791-020) cloning method.

All primers used in this study are listed in Supplementary Table 5.

### High-throughput sequencing

Bisulfite sequencing (BS-seq) libraries of meiocytes (of 18 different genotypes) and tapetal cells (two genotypes: WT and *rdr2*) were constructed as we previously described^24^, except for the incorporation of an extra round of bisulfite conversion. BS-seq libraries of the *pA9::RDR2 rdr2* and the *clsy3* mutant sperm were generated according to a previous description^11^. All BS-seq libraries were constructed from two biological replicates of isolated cells.

For siRNA-seq, total RNA was extracted from the isolated meiocytes (WT, *clsy3*, and *clsy4*) or tapetal cells using Direct-zol RNA Kit (Zymo Research, cat. no. R2061). Sequencing libraries were constructed using the RealSeq-Biofluids NGS Library Preparation Kit for miRNAs and small RNAs (BioCat, cat. no. 600-00048-SOM). Gel (Novex TBE 6%, Invitrogen, cat. no. EC6265BOX) selection and purification was performed to enrich libraries containing siRNA fragments ranging from 20 to 30 nt. Finally, the concentration and size distribution of purified siRNA libraries were analyzed using High Sensitivity DNA Chips (Agilent Technologies, cat. no. 5067-5594). All siRNA-seq libraries were constructed from two biological replicates of isolated cells.

RNA-seq libraries of pollen and tapetal cells of both WT and *drm* mutant backgrounds were produced as previously reported^11^. All RNA-seq libraries were constructed from three biological replicates of isolated cells.

All libraries were sequenced on NextSeq 500 (Illumina).

### CIRSPR/Cas9 deletion lines

CRISPR/Cas9 deletion constructs were generated using Golden Gate cloning into the *PICSL4723* destination vector according to a previous protocol^84^, but with additional single-guide RNAs (sgRNA; sequences listed in Supplementary Table 5). A vector containing 6 single-guide RNA cassettes was used to delete the HyperTE224 locus (HyperTEs are listed in Supplementary Table 2), and a vector containing 4 sgRNA cassettes (Supplementary Table 5) was created to delete HyperTE315 (Supplementary Table 2). Two independent T1 lines carrying 592-bp and 570-bp deletions at HyperTE224 were selected by Sanger sequencing (Supplementary Fig. 2a). Similarly, two T1 lines with 537-bp and 505-bp deletions at HyperTE315 were obtained (Supplementary Fig. 2c). For all four deletion lines, Cas9-free homozygous T2 plants were used for experiments.

### Immunolocalization

Inflorescences from *pA9::RDR2-Flag rdr2* plants were embedded in paraffin and sectioned transversely as previously described^33^. The dewaxed and rehydrated sections^33^ were treated with 1× PBS containing 100 μg/ml proteinase K for 10 minutes at room temperature, followed by 3 times of wash in 1× PBS. Next, the sections were incubated in 1× PBS containing 2.5% horse serum (Thermo Fisher Scientific, cat. no. 16050130) for 2h at room temperature and then rinsed with 1× PBS for 15 min. Subsequently, the sections were incubated in 1× PBS containing 1% (1:100 dilution) mouse monoclonal anti-Flag M2 primary antibody (Merck, cat. no. F1804) and 0.1% (w/v) BSA for 3-4h. Slides were then washed in 1× PBS twice for 10 minutes each and incubated for 2h with a goat anti-mouse Alexa Fluor 555 (Thermo Fisher Scientific, cat. no. A-21424) secondary antibody (1:500 dilution), followed by a wash for 10 min in 1× PBS. Finally, the sections were mounted under a coverslip with p-phenylenediamine anti-fade mounting agent^85^ and examined under a DM6000 microscope (Zeiss).

### Western blot

Inflorescences from *pA9::Pol-V-3*×*Flag pol-v* and *pA9::DCL3-3*×*Flag dcl3* were ground in liquid nitrogen and proteins were extracted and blotted as previously described^86^. For the primary antibody incubation, Anti-Flag (Merck, cat. no. F1804) was used at 1:2000 in 1× PBST solution dissolved with 5% skimmed milk. Blots were performed with the SuperSignal West Femto Maximum Sensitivity Substrate (Thermo Fisher Scientific, cat. no. 34095) and imaged using ImageQuant LAS500 (GE Healthcare).

### Confocal microscopy

Anthers were dissected from floral buds in a drop of sterilized water under a stereo microscope (Zeiss Stemi 1000) and staged as previously defined^87^. To examine whole seedlings, plants were grown on MS-agar plates with phosphinotricin (Melford, cat. no. 77182-82-2) for 10 days and mounted in a drop of sterilized water.

Dissected plant tissues or whole seedlings were visualized under a SP5 laser scanning confocal microscope (Leica Microsystems) using a 10× (HC PL FLUOTAR 10.0× 0.30 DRY) or 20× lens (HCX PL APO CS 20.0× 0.70 DRY UV). The following settings were used for the visualization of Venus (excitation 514 nm, emission band-pass at 511-563nm) and autofluorescence (excitation 514 nm, emission band-pass at 616-668 nm).

### Bisulfite-seq analysis

Low-quality sequencing reads, adaptor sequences and the first 9 bp of each read were removed using TrimGalore version 0.4.1 using default parameters. Filtered reads were then mapped to the TAIR10 *Arabidopsis* reference genome using Bismark version 0.22.2^88^. Duplicated reads were removed using Picard tools MarkDuplicates version 1.141. Subsequent DNA methylation analysis was performed as previously described^23^.

BS-seq data from seedlings^89^, cauline leaves^90^, rosette leaves^90,91^, WT and *rdr2* mutant roots^89,90^, WT and *drm* mutant meiocytes^11^, *rdr2* mutant sperm^11^, and WT^92^ and *drm* mutant sperm^22^ were obtained from published sources.

### siRNA-seq analysis

Adapter trimming for raw sequencing reads was performed using Cutadapt^93^. To determine HyperTEs, 21-nt to 25-nt meiotic siRNAs were mapped to the TAIR10 *Arabidopsis* reference genome with zero mismatches (-v 0) using Bowtie^94^ and clusters were determined with Shortstack^95^. Clusters were retained if they were above 99 bp and 24-nt log_2_(RPKM+1) > 7 and 24-nt log_2_(RPKM+1) > (25-nt log_2_(RPKM+1) + 2), resulting in 854 loci. These clusters were further filtered for significantly reduced CHH methylation (p < 0.01, Fisher’s Exact test) in *drm1 drm2* double mutant or *rdr2* mutant meiocytes compared to WT to obtain 797 clusters (HyperTEs, listed in Supplementary Table 2). To determine canonical RdDM (cRdDM) loci, previously defined somatic RdDM loci (9,993)^11^ were kept if they overlap neither sexual-lineage hypermethylated loci (1,257)^11^ nor the 797 HyperTEs, resulting in 9,618 loci. MetGenes were previously described (termed SLM, sexual-lineage-specific methylated loci)^11^.

Trimmed meiotic or tapetal 21-nt to 25-nt siRNA reads were mapped to the TAIR10 reference genome using Bowtie with either zero mismatches (-v 0) or up to three mismatches (-v 3). 24-nt siRNA abundance at each HyperTE, cRdDM, MetGene, gene, random control locus (determined previously^11^), and CLSY-dependent cluster^38^ was calculated using Reads Per Kilobase per Million (RPKM) of total mapped 24-nt siRNA reads or by total mapped 21-nt miRNA reads.

To identify connections between HyperTEs and MetGenes (Supplementary Table 3), 24-nt meiotic siRNAs were mapped to the TAIR10 genome with up to three mismatches (-v 3) using Bowtie. siRNAs overlapping MetGenes were retained and tagged with the MetGene location. These siRNAs were remapped to the TAIR10 genome a second time with zero mismatches (-v 0), and those mapped to HyperTEs were retained. Retained siRNAs mapped to the same MetGene and HyperTE were merged if they overlapped to form origin and target clusters. Abundance of zero mismatch meiotic and tapetal 24-nt siRNAs at each origin cluster was calculated using Reads Per Million (RPM) of total perfectly mapped 24-nt siRNA reads, and clusters were retained if RPM > 0 in both meiocyte and tapetum.

Besides our data, published sRNA data from *pol-iv* mutant meiocytes (Fig. 5)^96^, pollen^67^, and various tissues including floral buds^38^, seedlings^97,98^, leaves^96^ and roots^99^ were also used in this study.

### RNA-seq analysis

Transcription was calculated in FPKM (fragments per kilobase per million; Fig. 7a) as previous described^11^, and TPM (transcripts per million; Fig. 5b and Supplementary Table 4) using Kallisto^100^.

Published RNA-seq datasets for meiocytes and leaves from both WT and *drm* mutant backgrounds^11^ were also used in this study.

### Transposon and gene metaplots (ends analysis)

This was performed as described previously^23^.

### Plots and statistics

For all the violin plots in this study, the plot shows the distribution of the data and its probability density, with the box enclosing the middle 50% of the distribution (the horizontal line marks the median) and whiskers illustrating 1.5 times the interquartile range. Plots were generated for MetGenes (n= 469), HyperTEs (n= 797), cRdDMs (n= 9,618), or previously reported *clsy3*-(n= 804), *clsy4*-(n= 777) and *clsy1*,*2*-dependent (n=6,845) clusters^38^. All *P* values in violin plots were calculated by the Kolmogorov-Smirnov test. All reps included were biological replicates.

Scatter plots were generated from siRNA clusters as described in the corresponding figure legends with sample sizes and Pearson’s *R* values included.

### Imaging statistics

Confocal images of *GP1_LTR_::YFP drm/+* anthers (stage 8, containing microspores)^87^ were acquired with the settings for YFP and auto-fluorescence visualization as described in the Methods Details section. The fluorescent signals were recorded as images of 512 × 512 pixels, with a scan speed of 400Hz. All the images were acquired with the same SP5 setting for all the anthers. To quantify the microspores that are YFP positive, the image was inverted, converted into 8-bit, and switched to a binary mode using ImageJ^101^. The ‘watershed’ function was used to separate the neighboring microspores when the microspores were overlapped. The microspores with fluorescence were then automatically counted with the ‘analyze particles’ function. The total number of microspores with or without fluorescence was counted using the ‘point’ tool based on the images acquired in transmitted light (differential interference contrast, DIC) channel. Altogether, 41 anther locules from 13 different plants, containing 1391 microspores in total (705 microspores are YFP-positive, 50.7%), were analyzed.

## Acknowledgments

We thank the John Innes Centre Bioimaging Facility (Sergio Lopez, Eva Wegel and Kim Findlay) for their assistance with microscopy, and the Norwich BioScience Institute Partnership Computing infrastructure for Science Group for High Performance Computing resources. This work was funded by a European Research Council Starting Grant (‘SexMeth’ 804981; J.L., J.W., and X.F.), a Biotechnology and Biological Sciences Research Council (BBSRC) David Phillips Fellowship (BBL0250431; H.G. and X.F.), two BBSRC grants (BBS0096201 and BBP0135111; W.S. and X.F.), an EMBO Young Investigator Award (X.F.), a Sainsbury Charitable Foundation studentship (J.W.), and two John Innes Foundation studentships (B.A. and S.D.).

## Author contributions

J.L., J.W. and X.F. designed the study and wrote the manuscript, J.L., W.S., B.A., H.G., and S.D. performed the experiments, and J.L., J.W., B.A., H.G., and S.D. analyzed the data.

---

**Supplementary Fig. 1:**
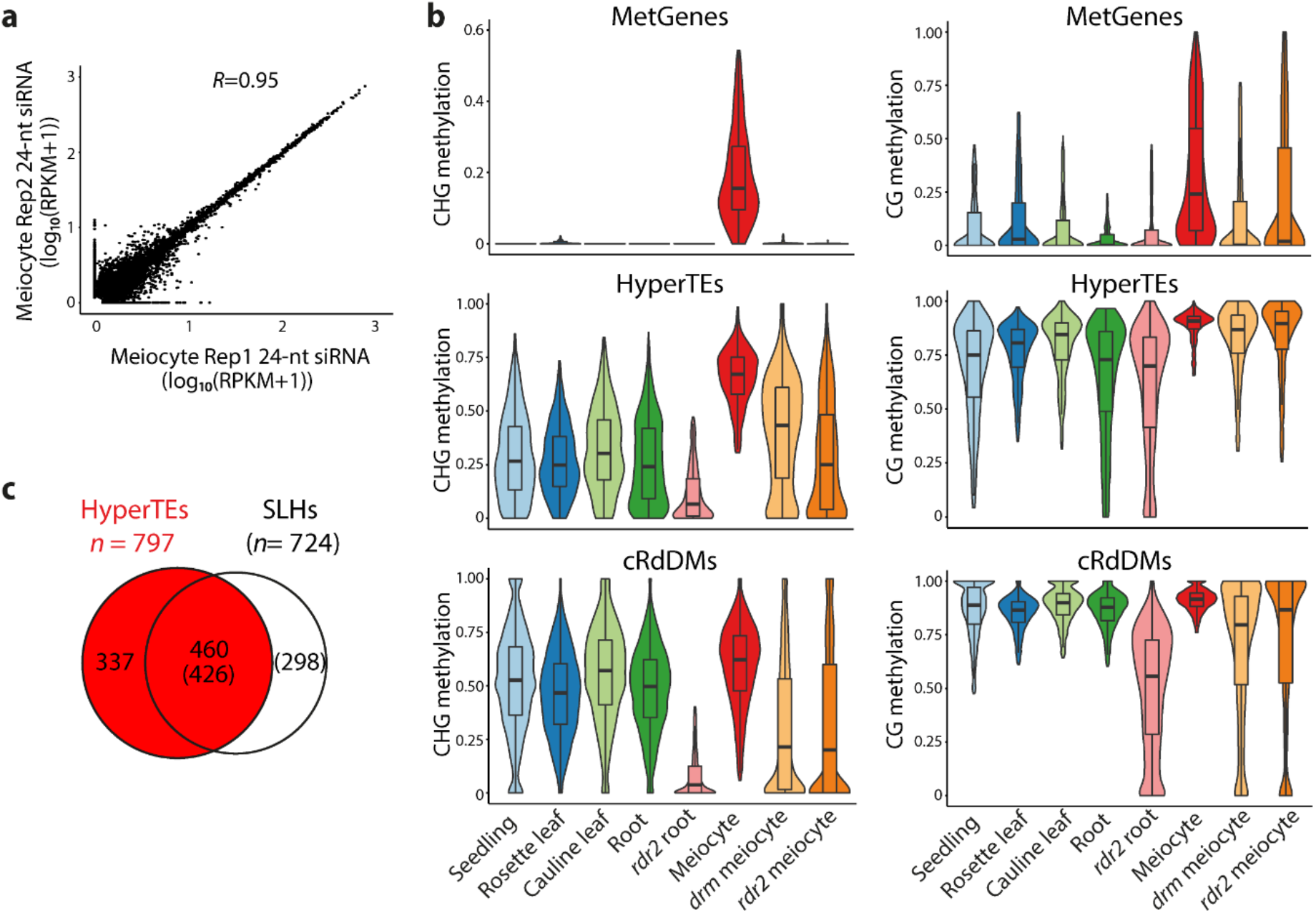
Male meiocytes have distinctive 24-nt siRNAs associated with HyperTEs. **a**, Scatterplot showing 24-nt siRNA abundance in the two biological replicates (Rep) of male meiocytes at identified clusters (n=4,822). Pearson’s *R*=0.95. **b**, Violin plots illustrating CHG methylation (left panels) and CG methylation (right panels) at MetGenes, HyperTEs, and canonical RdDM loci (cRdDMs) in somatic tissues and meiocytes. CHH methylation for these sites is shown in Fig. 1d. **c**, Scaled Venn diagram depicting the overlap between HyperTEs (797 loci) and published sexual-lineage hypermethylated loci (SLHs; 724 loci)^11^.

**Supplementary Fig. 2:**
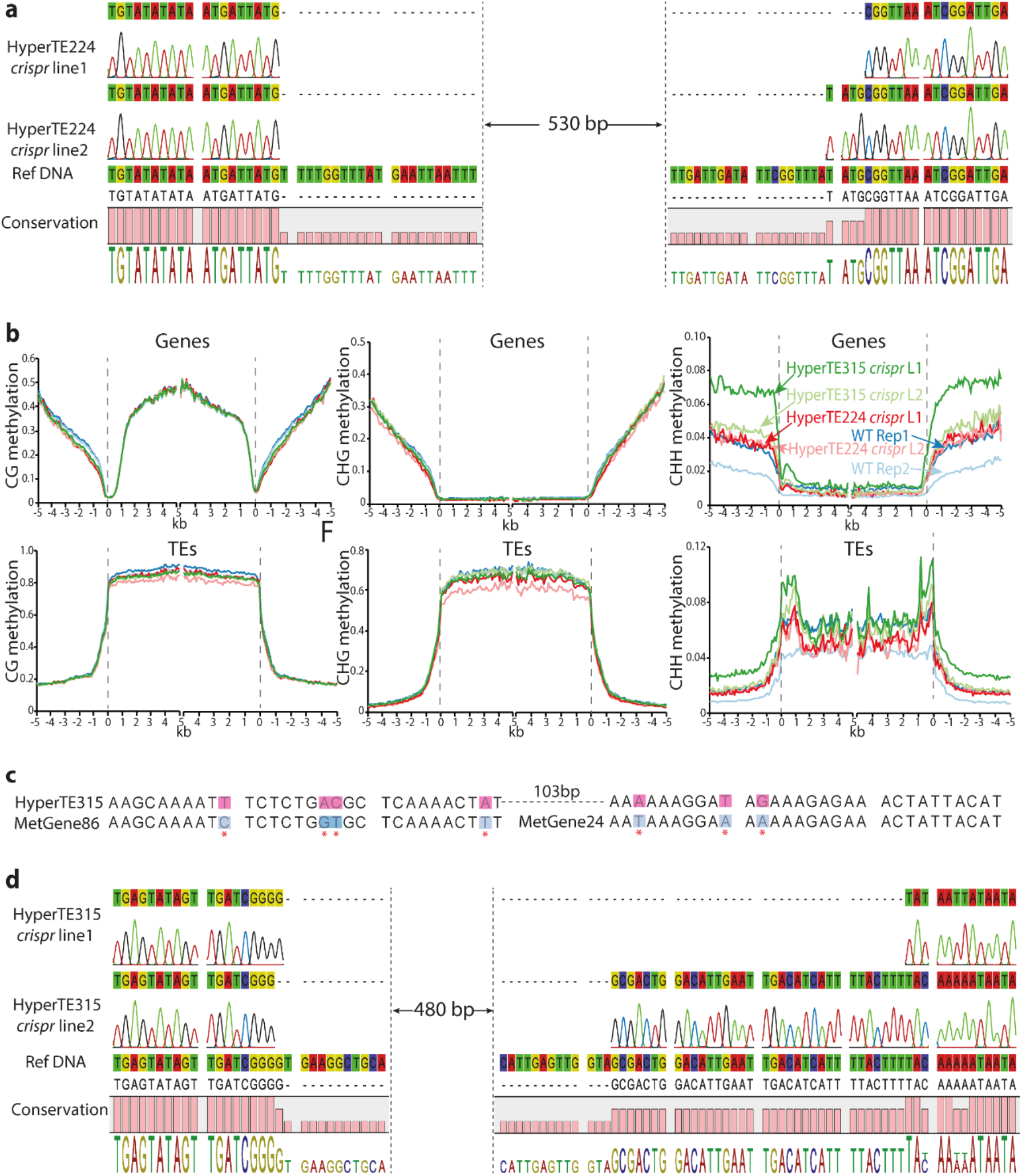
Overall methylation and sequences of HyperTE CRISPR deletion lines. **a**, **d**, DNA sequence alignment among two independent CRISPR/Cas9 deletion lines (**a** and **d**, respectively, for HyperTE224 and 315 deletion lines) and the reference genome (Ref DNA, TAIR 10). **b**, *Arabidopsis* genes or TEs aligned at the 5’ end (left subpanels) or the 3’ end (right subpanels) with average methylation in the CG, CHG or CHH context for each 100-bp interval plotted. The dashed line at zero represents the point of alignment. L, independent CRISPR line; Rep, biological replicate. **c**, DNA sequence alignment between HyperTE315 and the corresponding MetGene24 and MetGene86, as in Fig. 2b.

**Supplementary Fig. 3:**
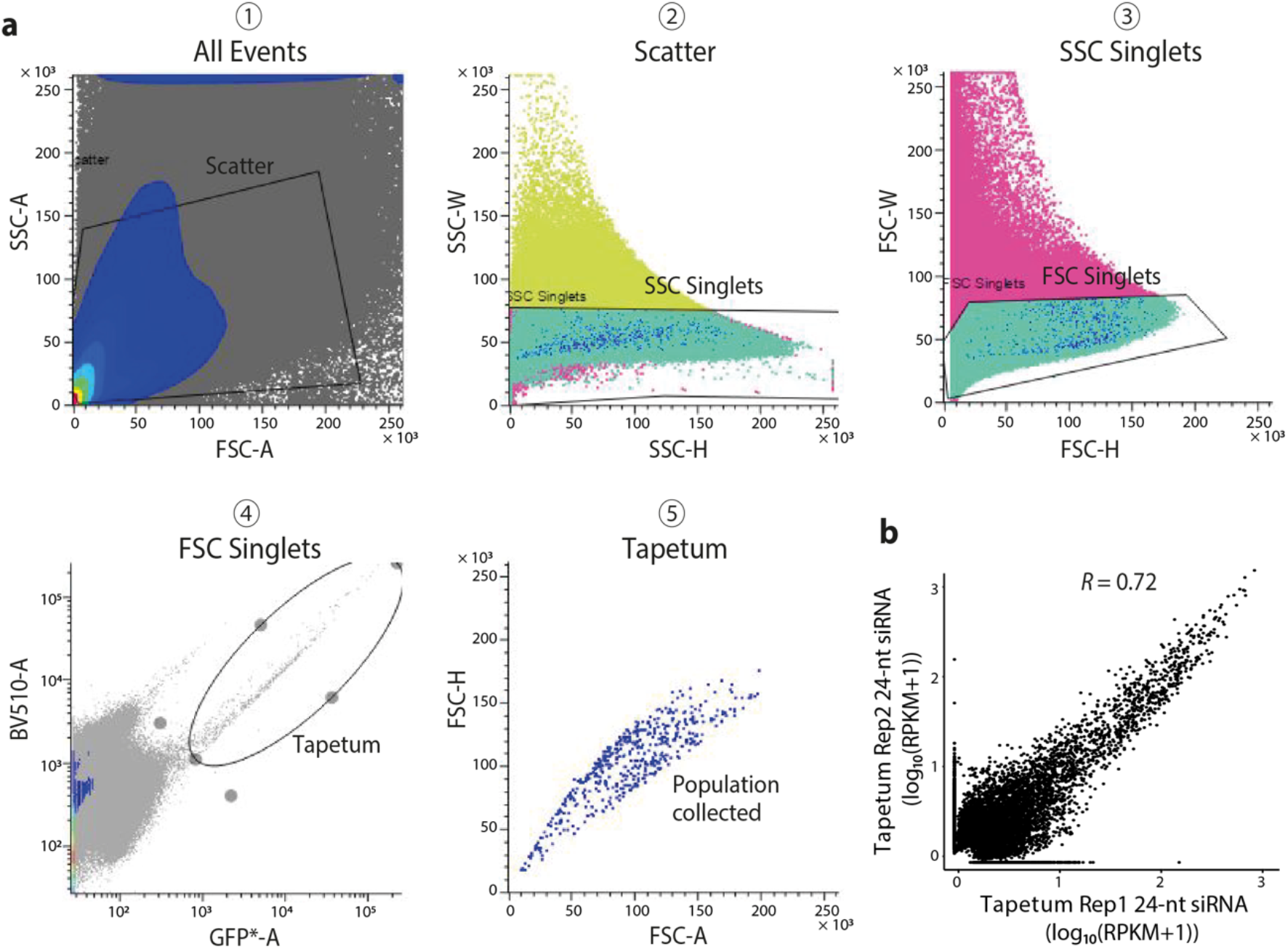
Tapetal cell isolation and siRNA-seq reproducibility. **a**, Gate settings used in the FACS of tapetal cells from *pTP::GFP* plants. **b**, Scatterplot showing 24-nt siRNA abundance in the two biological replicates of tapetal cells at identified clusters (n=8,476), as in Supplementary Fig. 1a. Pearson’s *R*=0.72.

**Supplementary Fig. 4:**
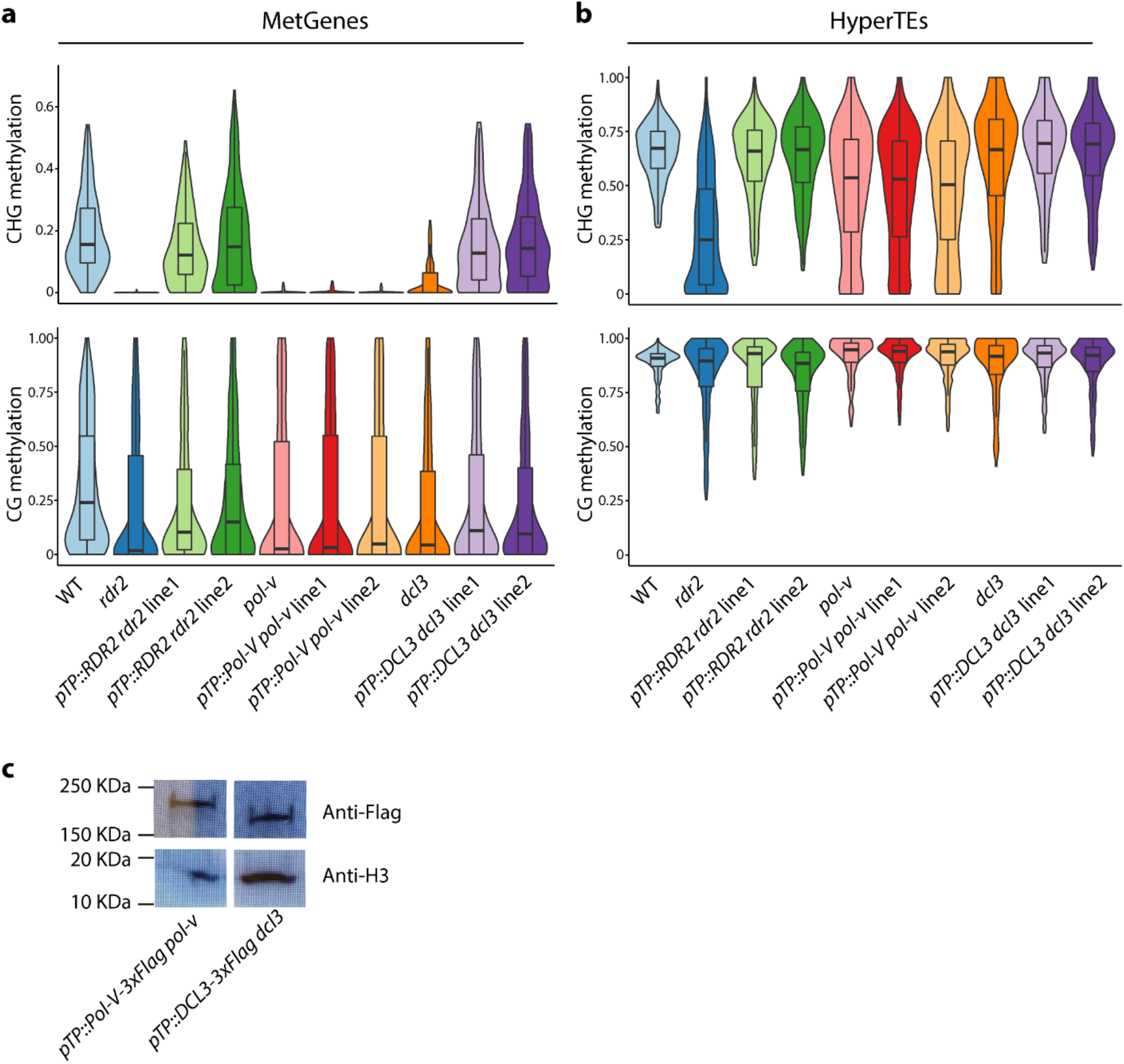
Tapetal 24-nt siRNAs are sufficient to drive methylation reprogramming in meiocytes. **a**, **b**, Violin plots depicting CHG (upper panels) and CG methylation (lower panels) at MetGenes (**a**) and HyperTEs (**b**) in the meiocytes (MC) from different genotypes. CHH methylation illustrated in Fig. 4b. **c**, Western blot showing Pol V-3xFlag and DCL3-3xFlag fusion proteins detected from *pTP::Pol-V pol-v* and *pTP::DCL3 dcl3* young flower buds, respectively. Anti-H3 antibody was used as a control.

**Supplementary Fig. 5:**
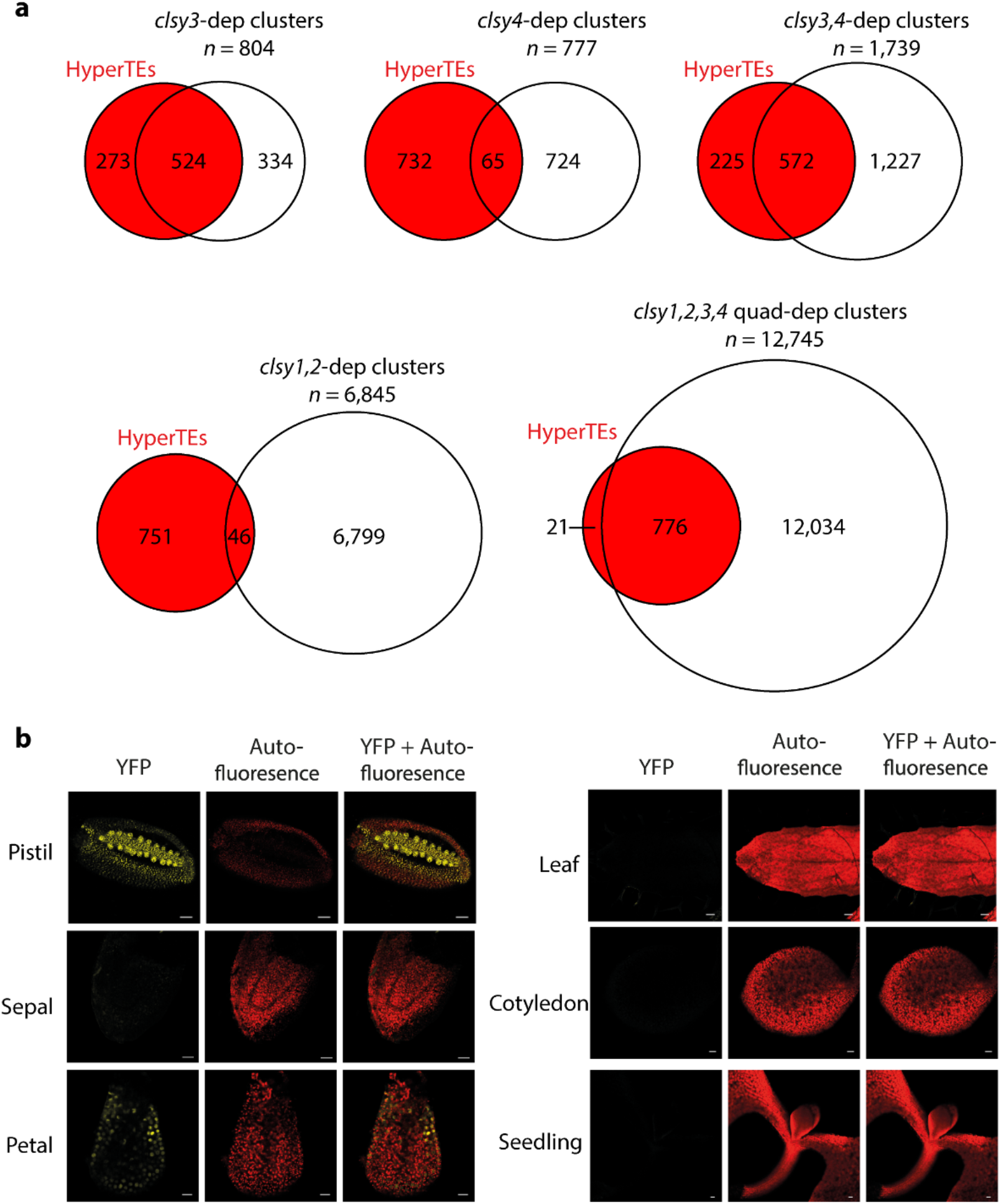
HyperTEs largely overlap siRNA clusters that require CLSY3, which shows tissue-specific localization. **a**, Scaled Venn diagrams illustrating the overlap between HyperTEs (797 loci) and published *clsy*-dependent clusters. **b**, Confocal microscopy images showing CLSY3-Venus localization in different *Arabidopsis* tissues. YFP signal was observed in pistils and petals. Yellow, YFP; red, autofluorescence. Scale bars: pistil, 25 μm; sepal, 30 μm; petal, 10 μm; leaf, 100 μm; cotyledon and seedling, 50 μm.

**Supplementary Fig. 6:**
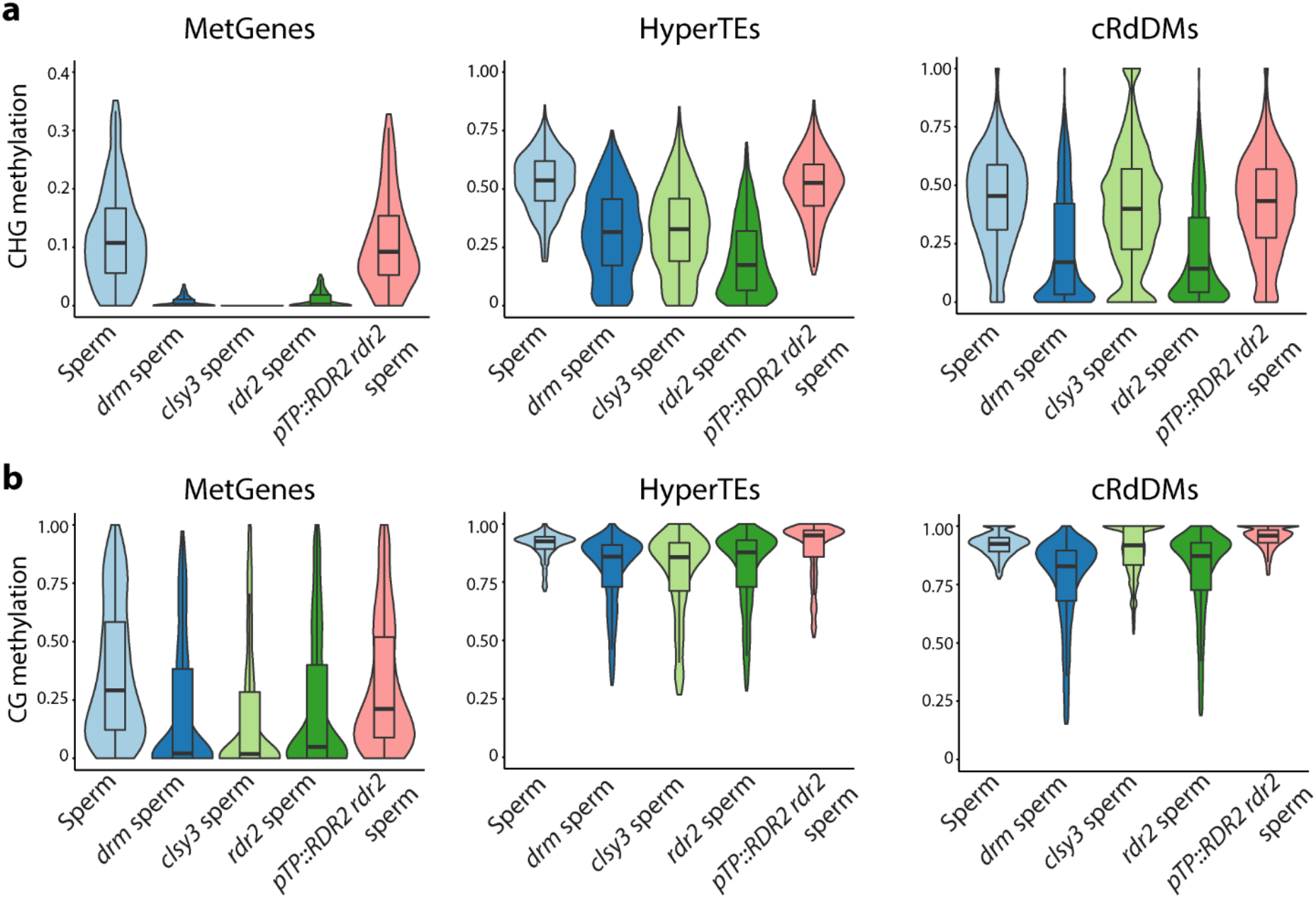
Tapetal niRNAs are sufficient to reconstitute sperm DNA methylation. Violin plots presenting CHG (**a**) and CG (**b**) methylation at MetGenes, HyperTEs and cRdDMs in the sperm from WT and different genotypes. CHH methylation is shown in Fig. 6a.

**Supplementary Fig. 7:**
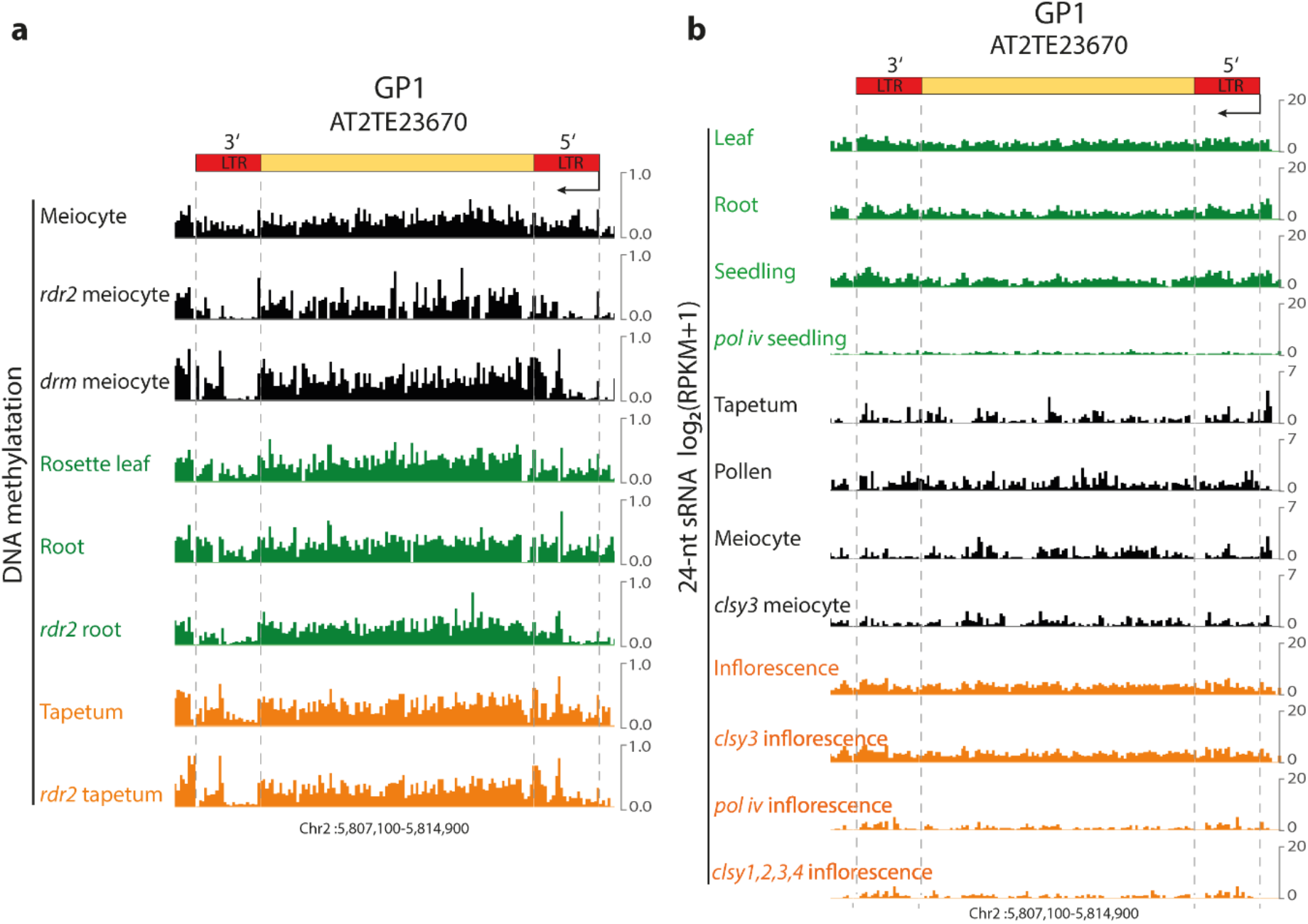
DNA methylation and siRNA abundance at the *GP1* transposon. **a**, **b**, Snapshots showing DNA methylation (**a**) and siRNA abundance (**b**) at *GP1* in WT and RdDM mutant cells and tissues. LTRs of *GP1* lose substantial methylation in RdDM mutants in meiocytes, tapetal cells and soma. Consistently, 24-nt siRNAs at *GP1* are greatly reduced in the *pol-iv* and the *clsy1,2,3,4* quadruple mutant, although not in the *clsy3* single mutant.

## References

1. Smith, Z.D. & Meissner, A. DNA methylation: roles in mammalian development. Nat Rev Genet 14, 204–20 (2013).

2. Zhang, H., Lang, Z. & Zhu, J.K. Dynamics and function of DNA methylation in plants. Nat Rev Mol Cell Biol 19, 489–506 (2018).

3. Zemach, A. & Zilberman, D. Evolution of eukaryotic DNA methylation and the pursuit of safer sex. Curr Biol 20, R780–5 (2010).

4. Law, J.A. & Jacobsen, S.E. Establishing, maintaining and modifying DNA methylation patterns in plants and animals. Nat Rev Genet 11, 204–20 (2010).

5. Matzke, M.A. & Mosher, R.A. RNA-directed DNA methylation: an epigenetic pathway of increasing complexity. Nat Rev Genet 15, 394–408 (2014).

6. Wendte, J.M. & Pikaard, C.S. The RNAs of RNA-directed DNA methylation. Biochim Biophys Acta Gene Regul Mech 1860, 140–148 (2017).

7. Kuo, H.Y., Jacobsen, E.L., Long, Y., Chen, X. & Zhai, J. Characteristics and processing of Pol IV-dependent transcripts in Arabidopsis. J Genet Genomics 44, 3–6 (2017).

8. Pikaard, C.S. & Mittelsten Scheid, O. Epigenetic regulation in plants. Cold Spring Harb Perspect Biol 6, a019315 (2014).

9. Seisenberger, S. et al. Reprogramming DNA methylation in the mammalian life cycle: building and breaking epigenetic barriers. Philosophical Transactions of the Royal Society B-Biological Sciences 368 (2013).

10. Tang, W.W.C., Kobayashi, T., Irie, N., Dietmann, S. & Surani, M.A. Specification and epigenetic programming of the human germ line. Nature Reviews Genetics 17, 585–600 (2016).

11. Walker, J. et al. Sexual-lineage-specific DNA methylation regulates meiosis in Arabidopsis. Nature Genetics 50, 130–137 (2018).

12. Toth, K.F., Pezic, D., Stuwe, E. & Webster, A. The piRNA pathway guards the germline genome against transposable elements. Adv Exp Med Biol 886, 51–77 (2016).

13. Greenberg, M.V.C. & Bourc’his, D. The diverse roles of DNA methylation in mammalian development and disease. Nature Reviews Molecular Cell Biology 20, 590–607 (2019).

14. Stewart, K.R., Veselovska, L. & Kelsey, G. Establishment and functions of DNA methylation in the germline. Epigenomics 8, 1399–1413 (2016).

15. Parfrey, L.W., Lahr, D.J.G., Knoll, A.H. & Katz, L.A. Estimating the timing of early eukaryotic diversification with multigene molecular clocks. Proceedings of the National Academy of Sciences of the United States of America 108, 13624–13629 (2011).

16. Schmidt, A., Schmid, M.W. & Grossniklaus, U. Plant germline formation: common concepts and developmental flexibility in sexual and asexual reproduction. Development 142, 229–41 (2015).

17. Hackenberg, D. & Twell, D. The evolution and patterning of male gametophyte development. Plant Development and Evolution 131, 257–298 (2019).

18. Vielle-Calzada, J.P. Linking stem cells to germ cells. Science 356, 378–379 (2017).

19. Feng, X., Zilberman, D. & Dickinson, H. A conversation across generations: soma-germ cell crosstalk in plants. Dev Cell 24, 215–25 (2013).

20. Gomez, J.F., Talle, B. & Wilson, Z.A. Anther and pollen development: A conserved developmental pathway. J Integr Plant Biol 57, 876–91 (2015).

21. Quadrana, L. & Colot, V. Plant Transgenerational Epigenetics. Annual Review of Genetics, Vol 50 50, 467–491 (2016).

22. Hsieh, P.H. et al. Arabidopsis male sexual lineage exhibits more robust maintenance of CG methylation than somatic tissues. Proc Natl Acad Sci U S A 113, 15132–15137 (2016).

23. Ibarra, C.A. et al. Active DNA demethylation in plant companion cells reinforces transposon methylation in gametes. Science 337, 1360–4 (2012).

24. Park, K. et al. DNA demethylation is initiated in the central cells of Arabidopsis and rice. Proc Natl Acad Sci U S A 113, 15138–15143 (2016).

25. Calarco, J.P. et al. Reprogramming of DNA methylation in pollen guides epigenetic inheritance via small RNA. Cell 151, 194–205 (2012).

26. Mamun, E.A., Cantrill, L.C., Overall, R.L. & Sutton, B.G. Cellular organisation and differentiation of organelles in pre-meiotic rice anthers. Cell Biol Int 29, 792–802 (2005).

27. Sager, R. & Lee, J.Y. Plasmodesmata in integrated cell signalling: insights from development and environmental signals and stresses. J Exp Bot 65, 6337–58 (2014).

28. Melnyk, C.W., Molnar, A. & Baulcombe, D.C. Intercellular and systemic movement of RNA silencing signals. EMBO J 30, 3553–63 (2011).

29. Pyott, D.E. & Molnar, A. Going mobile: non-cell-autonomous small RNAs shape the genetic landscape of plants. Plant Biotechnol J 13, 306–18 (2015).

30. Liu, L. & Chen, X. Intercellular and systemic trafficking of RNAs in plants. Nat Plants 4, 869–878 (2018).

31. Zhai, J. et al. Spatiotemporally dynamic, cell-type-dependent premeiotic and meiotic phasiRNAs in maize anthers. Proc Natl Acad Sci U S A 112, 3146–51 (2015).

32. Paul, W., Hodge, R., Smartt, S., Draper, J. & Scott, R. The isolation and characterization of the tapetum-specific Arabidopsis thaliana A9 gene. Plant Molecular Biology 19, 611–622 (1992).

33. Feng, X. & Dickinson, H.G. Tapetal cell fate, lineage and proliferation in the Arabidopsis anther. Development 137, 2409–16 (2010).

34. Turgut, K. et al. The highly expressed tapetum-specific A9 gene is not required for male fertility in Brassica napus. Plant Molecular Biology 24, 97–104 (1994).

35. Lee, Y.H. et al. Induction of male sterile cabbage using a tapetum-specific promoter from Brassica campestris L. ssp pekinensis. Plant Cell Reports 22, 268–273 (2003).

36. Xu, X.F. et al. Magnesium Transporter 5 plays an important role in Mg transport for male gametophyte development in Arabidopsis. Plant J 84, 925–36 (2015).

37. Faulkner, C. Plasmodesmata and the symplast. Current Biology 28, R1374–R1378 (2018).

38. Zhou, M., Palanca, A.M.S. & Law, J.A. Locus-specific control of the de novo DNA methylation pathway in Arabidopsis by the CLASSY family. Nature Genetics 50, 865–873 (2018).

39. Gallego-Bartolome, J. et al. Co-targeting RNA Polymerases IV and V Promotes Efficient De Novo DNA Methylation in Arabidopsis. Cell 176, 1068–1082 (2019).

40. Smith, L.M. et al. An SNF2 protein associated with nuclear RNA silencing and the spread of a silencing signal between cells in Arabidopsis. Plant Cell 19, 1507–1521 (2007).

41. Dunoyer, P., Himber, C., Ruiz-Ferrer, V., Alioua, A. & Voinnet, O. Intra- and intercellular RNA interference in Arabidopsis thaliana requires components of the microRNA and heterochromatic silencing pathways. Nat Genet 39, 848–56 (2007).

42. Lewsey, M.G. et al. Mobile small RNAs regulate genome-wide DNA methylation. Proc Natl Acad Sci U S A 113, E801–10 (2016).

43. Molnar, A. et al. Small silencing RNAs in plants are mobile and direct epigenetic modification in recipient cells. Science 328, 872–5 (2010).

44. Melnyk, C.W., Molnar, A., Bassett, A. & Baulcombe, D.C. Mobile 24 nt small RNAs direct transcriptional gene silencing in the root meristems of Arabidopsis thaliana. Curr Biol 21, 1678–83 (2011).

45. Slotkin, R.K. et al. Epigenetic reprogramming and small RNA silencing of transposable elements in pollen. Cell 136, 461–472 (2009).

46. Fei, Q.L., Yang, L., Liang, W.Q., Zhang, D.B. & Meyers, B.C. Dynamic changes of small RNAs in rice spikelet development reveal specialized reproductive phasiRNA pathways. Journal of Experimental Botany 67, 6037–6049 (2016).

47. Kakrana, A. et al. Plant 24-nt reproductive phasiRNAs from intramolecular duplex mRNAs in diverse monocots. Genome Research 28, 1333–1344 (2018).

48. Xia, R. et al. 24-nt reproductive phasiRNAs are broadly present in angiosperms. Nature Communications 10 (2019).

49. Ono, S. et al. EAT1 transcription factor, a non-cell-autonomous regulator of pollen production, activates meiotic small RNA biogenesis in rice anther tapetum. Plos Genetics 14 (2018).

50. Song, X.W. et al. Roles of DCL4 and DCL3b in rice phased small RNA biogenesis. Plant Journal 69, 462–474 (2012).

51. Teng, C. et al. Dicer-like 5 deficiency confers temperature-sensitive male sterility in maize. Nature Communications 11 (2020).

52. Johnson, C. et al. Clusters and superclusters of phased small RNAs in the developing inflorescence of rice. Genome Research 19, 1429–1440 (2009).

53. Komiya, R. et al. Rice germline-specific Argonaute MEL1 protein binds to phasiRNAs generated from more than 700 lincRNAs. Plant J 78, 385–97 (2014).

54. Nonomura, K.I. et al. A germ cell-specific gene of the ARGONAUTE family is essential for the progression of premeiotic mitosis and meiosis during sporogenesis in rice. Plant Cell 19, 2583–2594 (2007).

55. Dukowic-Schulze, S. et al. Novel meiotic miRNAs and indications for a role of phasiRNAs in meiosis. Frontiers in Plant Science 7 (2016).

56. Zhang, M. et al. CHH DNA methylation increases at 24-PHAS loci depend on 24-nt phasiRNAs in maize meiotic anthers. New Phytol (2020).

57. Ozata, D.M., Gainetdinov, I., Zoch, A., O’Carroll, D. & Zamore, P.D. PIWI-interacting RNAs: small RNAs with big functions. Nature Reviews Genetics 20, 89–108 (2019).

58. Borges, F. et al. Transposon-derived small RNAs triggered by miR845 mediate genome dosage response in Arabidopsis. Nat Genet 50, 186–192 (2018).

59. Martinez, G. et al. Paternal easiRNAs regulate parental genome dosage in Arabidopsis. Nat Genet 50, 193–198 (2018).

60. Martinez, G. & Kohler, C. Role of small RNAs in epigenetic reprogramming during plant sexual reproduction. Curr Opin Plant Biol 36, 22–28 (2017).

61. Wang, G.F., Jiang, H., de Leon, G.D., Martinez, G. & Kohler, C. Sequestration of a transposon-derived siRNA by a target mimic imprinted gene induces postzygotic reproductive isolation in Arabidopsis. Developmental Cell 46, 696–705 (2018).

62. Yu, Y., Zhang, Y.C., Chen, X.M. & Chen, Y.Q. Plant noncoding RNAs: hidden players in development and stress responses. Annual Review of Cell and Developmental Biology 35, 407–431 (2019).

63. Su, Z.X. et al. Regulation of female germline specification via small RNA mobility in Arabidopsis. Plant Cell 32, 2842–2854 (2020).

64. Kirkbride, R.C. et al. Maternal small RNAs mediate spatial-temporal regulation of gene expression, imprinting, and seed development in Arabidopsis. Proceedings of the National Academy of Sciences of the United States of America 116, 2761–2766 (2019).

65. Olmedo-Monfil, V. et al. Control of female gamete formation by a small RNA pathway in Arabidopsis. Nature 464, 628–632 (2010).

66. Sarkies, P. et al. Ancient and novel small RNA pathways compensate for the loss of piRNAs in multiple independent nematode lineages. Plos Biology 13 (2015).

67. Martinez, G., Panda, K., Kohler, C. & Slotkin, R.K. Silencing in sperm cells is directed by RNA movement from the surrounding nurse cell. Nat Plants 2, 16030 (2016).

68. Grant-Downton, R. et al. Artificial microRNAs reveal cell-specific differences in small RNA activity in pollen. Curr Biol 23, R599–601 (2013).

69. Kawashima, T. & Berger, F. Epigenetic reprogramming in plant sexual reproduction. Nat Rev Genet 15, 613–24 (2014).

70. Chow, H.T., Chakraborty, T. & Mosher, R.A. RNA-directed DNA Methylation and sexual reproduction: expanding beyond the seed. Current Opinion in Plant Biology 54, 11–17 (2020).

71. Wang, G.F. & Kohler, C. Epigenetic processes in flowering plant reproduction. Journal of Experimental Botany 68, 797–807 (2017).

72. Kiuchi, T. et al. A single female-specific piRNA is the primary determiner of sex in the silkworm. Nature 509, 633–6 (2014).

73. Ernst, C., Odom, D.T. & Kutter, C. The emergence of piRNAs against transposon invasion to preserve mammalian genome integrity. Nature Communications 8 (2017).

74. Rojas-Rios, P. & Simonelig, M. piRNAs and PIWI proteins: regulators of gene expression in development and stem cells. Development 145 (2018).

75. Dai, P. et al. A translation-activating function of MIWI/piRNA during mouse spermiogenesis. Cell 179, 1566–1581 (2019).

## References

76. Xie, Z. et al. Genetic and functional diversification of small RNA pathways in plants. PLoS Biol 2, E104 (2004).

77. Pontier, D. et al. Reinforcement of silencing at transposons and highly repeated sequences requires the concerted action of two distinct RNA polymerases IV in Arabidopsis. Genes & Development 19, 2030–2040 (2005).

78. Blevins, T. et al. Four plant Dicers mediate viral small RNA biogenesis and DNA virus induced silencing. Nucleic Acids Res 34, 6233–46 (2006).

79. Cao, X. & Jacobsen, S.E. Role of the arabidopsis DRM methyltransferases in De novo DNA methylation and Gene silencing Curr Biol 12, 1138–44 (2002).

80. Park, K. et al. Optimized methods for the isolation of Arabidopsis female central cells and their nuclei. Molecules and Cells 39, 768–775 (2016).

81. Deal, R.B. & Henikoff, S. The INTACT method for cell type-specific gene expression and chromatin profiling in Arabidopsis thaliana. Nat Protoc 6, 56–68 (2011).

82. Nakagawa, T. et al. Development of series of gateway binary vectors, pGWBs, for realizing efficient construction of fusion genes for plant transformation. J Biosci Bioeng 104, 34–41 (2007).

83. Karimi, M., De Meyer, B. & Hilson, P. Modular cloning in plant cells. Trends in Plant Science 10, 103–105 (2005).

84. Castel, B., Tomlinson, L., Locci, F., Yang, Y. & Jones, J.D.G. Optimization of T-DNA architecture for Cas9-mediated mutagenesis in Arabidopsis. PLoS One 14, e0204778 (2019).

85. Krenik, K.D., Kephart, G.M., Offord, K.P., Dunnette, S.L. & Gleich, G.J. Comparison of antifading agents used in immunofluorescence. Journal of Immunological Methods 117, 91–97 (1989).

86. Yang, H. et al. Genome-Wide Mapping of Targets of Maize Histone Deacetylase HDA101 Reveals Its Function and Regulatory Mechanism during Seed Development. Plant Cell 28, 629–45 (2016).

87. Sanders, P.M. et al. Anther developmental defects in Arabidopsis thaliana male-sterile mutants. Sexual Plant Reproduction 11, 297–322 (1999).

88. Krueger, F. & Andrews, S.R. Bismark: a flexible aligner and methylation caller for Bisulfite-Seq applications. Bioinformatics 27, 1571–1572 (2011).

89. Zemach, A. et al. The Arabidopsis nucleosome remodeler DDM1 allows DNA methyltransferases to access H1-containing heterochromatin. Cell 153, 193–205 (2013).

90. Coleman-Derr, D. & Zilberman, D. Deposition of histone variant H2A.Z within gene bodies regulates responsive genes. PLoS Genet 8, e1002988 (2012).

91. Stroud, H. et al. Non-CG methylation patterns shape the epigenetic landscape in Arabidopsis. Nat Struct Mol Biol 21, 64–72 (2014).

92. He, S.B., Vickers, M., Zhang, J.Y. & Feng, X.Q. Natural depletion of histone H1 in sex cells causes DNA demethylation, heterochromatin decondensation and transposon activation. Elife 8, e42530 (2019).

93. Martin, M. Cutadapt removes adapter sequences from high-throughput sequencing reads. EMBnet. journal 17, 10–12 (2011).

94. Langmead, B. Aligning short sequencing reads with Bowtie. Curr Protoc Bioinformatics 11, 7 (2010).

95. Johnson, N.R., Yeoh, J.M., Coruh, C. & Axtell, M.J. Improved Placement of Multi-mapping Small RNAs. G3-Genes Genomes Genetics 6, 2103–2111 (2016).

96. Huang, J.Y. et al. Meiocyte-specific and AtSPO11-1-dependent small RNAs and their association with meiotic gene expression and recombination. Plant Cell 31, 444–464 (2019).

97. Schalk, C. et al. Small RNA-mediated repair of UV-induced DNA lesions by the DNA DAMAGE-BINDING PROTEIN 2 and ARGONAUTE 1. Proceedings of the National Academy of Sciences of the United States of America 114, E2965–E2974 (2017).

98. Ye, R.Q. et al. A Dicer-independent route for biogenesis of siRNAs that direct DNA methylation in Arabidopsis. Molecular Cell 61, 222–235 (2016).

99. Vidal, E.A. et al. Integrated RNA-seq and sRNA-seq analysis identifies novel nitrate-responsive genes in Arabidopsis thaliana roots. Bmc Genomics 14 (2013).

100. Bray, N.L., Pimentel, H., Melsted, P. & Pachter, L. Near-optimal probabilistic RNA-seq quantification. Nature Biotechnology 34, 525–527 (2016).

101. Schneider, C.A., Rasband, W.S. & Eliceiri, K.W. NIH Image to ImageJ: 25 years of image analysis. Nat Methods 9, 671–5 (2012).

